# Thermodynamic Insights into the WT and Y220C TP53 DBDs Reveals that the Oncogenic Y220C Variant is a Loss of Function Mutation for Zn^2+^-binding at Physiological Temperature

**DOI:** 10.64898/2026.01.31.701171

**Authors:** Jacob I. Stuckey, Valerie Vivat, Bradley M. Dickson, Jeremy W. Setser, Yong Li, Claudia M. Cremers, Jonathan E. Wilson, Daniele Peterle, John R. Engen, Thomas E. Wales, Rainer Wilcken, Erin B. Rohan, Gregg Chenail, James E. Audia, Robert J. Sims

## Abstract

We present evidence of previously unrecognized allosteric connectivity across the TP53 DNA binding domain (DBD). Specifically, we have found evidence of explicit influence on the Zn^2+^-binding site from the region surrounding the hotspot Y220C mutation. This allosteric connectivity is intertwined with a temperature-dependent destabilization of Zn^2+^ binding in both the WT and Y220C DBDs. Our studies indicate that the Y220C mutation exacerbates this temperature-dependent destabilization of Zn^2+^-binding to result in overall destabilization of the Y220C variant. We provide detailed thermodynamic evidence that Rezatapopt, a small molecule reactivator of the Y220C DBD, engages Y220C through two distinct thermodynamic pathways and restores WT-level Zn^2+^-affinity to this oncogenic variant. A series of thermodynamic models describing the WT and Y220C conformational landscapes, as well as the Rezatapopt binding mechanisms, are proposed.

## Introduction

Mutations in the *TP53* tumor suppressor gene are among the most common in human cancers with ∼50% of tumors showing alteration in *TP53*. The Y220C mutation is the most prevalent mutation away from the DNA-binding surface. This mutation creates a solvent accessible cleft at the periphery of the β-Sandwich at the core of the DBD^1^. This cleft has been thoroughly explored for its small molecule binding capacity with robust evidence of noncovalent targeting, as well as evidence of direct covalent targeting of the mutant cysteine residue. The most advanced of any Y220C-targeting molecule is Rezatapopt, a noncovalent, reversible reactivator of Y220C currently in clinical development^2–8^.

Y220C was originally classified as a thermally destabilizing mutation of the TP53 DNA Binding Domain (DBD)^9^. This contrasts with one of the most prevalent TP53 mutations, R175H, shown to weaken the DBD affinity for zinc. It is worth noting that one study of Y220C acknowledged that it could not exclude the possibility that this mutation has not been properly categorized and could actually be a mixed zinc-binding/stability class mutation^10^.

All studies identified in our sweep of the literature do not actually utilize the endogenous Y220C DBD but rather a “superstable quadruple mutant” to increase the stability of the construct (M133L/V203A/N239Y/N268D)^11^. Further, we identified one report that extension of the TP53 wild-type (WT) DBD by 5 additional N-terminal residues beyond its traditional domain boundaries conferred increased thermal stability and a decreased propensity for aggregation^12^. Despite this report, there seems to have been minimal uptake of this N-terminally extended construct for use in studying the TP53 DBD, particularly the Y220C variant^2–4,7,10^.

We therefore set out to study the endogenous Y220C DBD sequence to enable us to bypass potential confounding influences of stabilizing alterations. We further explored extension of the N-terminal boundary to examine its influence. This improved protein construct design allowed us to better understand the nature of the Y220C mutation, affording us insights into the mechanism of increased stability of the Y220C variant by small molecule reactivators. Comparison of Y220C and WT behavior across a suite of assays also revealed novel, fundamental insights into the conformational landscape of the WT DBD as well. A conserved water network has been identified previously in the region of the protein around residue 220^13^ and this network integrates seamlessly with a holistic interpretation of our experimental observations.

## Results

### Protein Construct Design and Characterization

To study the Y220C DBD, we were able to express the endogenous TP53 Y220C DBD without stabilizing mutations using the traditional boundary (94-312), but our focus was on expression of a TP53 Y220C DBD using, not only the endogenous sequence, but also beginning at residue 89 ^12^. Thorough biophysical characterization of this construct inspired a C-terminally truncated construct utilized for crystallographic studies (residues 89-293). In short, we observed overall consistent structuring of these constructs with the WT DBD and an increased thermal stability of 1.4°C relative to the traditional, truncated N-terminal boundary. Additionally, we observed a 7.87°C thermal destabilization of this endogenous mutant construct relative to boundary-matched WT (**Supplementary Figure 1**).

### Differential Rescue by Rezatapopt with the Extended Y220C DBD Relative to the Truncated Y220C DBD Suggests Allosteric Connectivity between the Y220C Pocket and the Zn^2+^-Coordination Site

We sought to understand differences in the ability of small molecule correctors to stabilize the TP53 Y220C truncated DBD (94-312) vs. the N-terminally extended TP53 Y220C DBD (89-312). To do this, we utilized Rezatapopt for its ability to rescue DNA binding upon thermal challenge of the respective domains (**Fig. 1a**). We compared each construct to a boundary-matched, WT version. Briefly, constructs were heated to 28°C for 30 minutes in the presence or absence of compound, analogous to previous characterization efforts^4^. Samples were cooled to room temperature and residual DNA binding capacity and affinity were compared to the WT, boundary-matched construct using a TR-FRET-based DNA binding assay.

**Figure 1.**
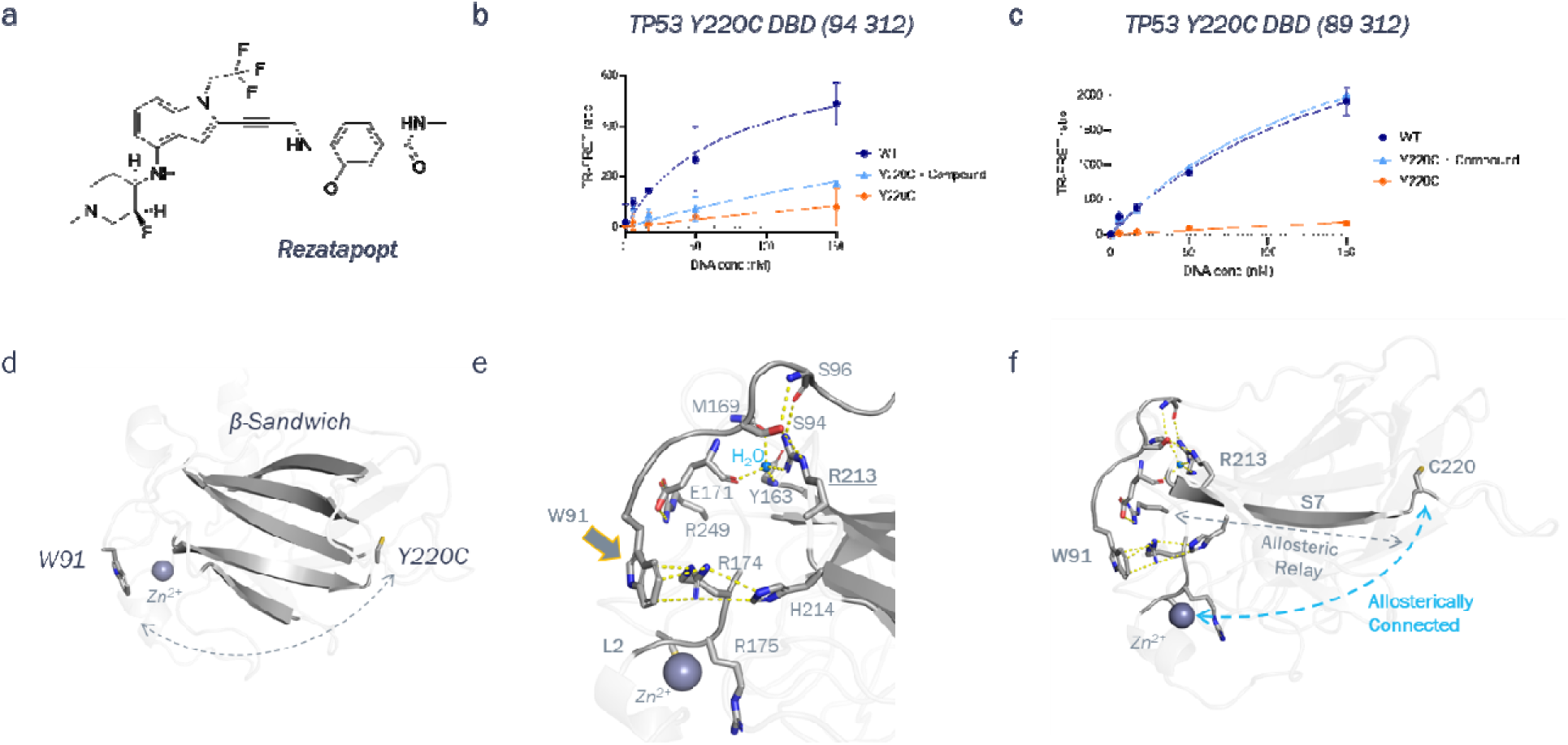
Proposed Allosteric Connectivity between Y220C and Zn^2+^ inspired by compound studies with traditional and extended DBD boundaries. (**a**) Structure of Rezatapopt. (**b**) Residual DNA binding affinity after heat challenge at 30°C with truncated TP53 Y220C DBD (±) 10 µM Rezatapopt. WT control is boundary and sequence-matched. (**c**) Residual DNA binding affinity after heat challenge at 30°C for N-terminally extended, endogenous Y220C DBD (±) 10 µM Rezatapopt. WT control is boundary and sequence-matched. For (**b**) and (**c**), experiments were performed in duplicate and error bars represent the standard error of the mean. (**d**) Spatial disparity between Rezatapopt binding pocket/Y220C site and W91. (**e**) Examination of W91 proximity and participation in an intricate structural network near key features of Zn^2+^-coordination site (**f**) Proposed allosteric connectivity from Y220C to Zn^2+^-coordination site mediated through S7 β-strand and the intricate structural network the S7 strand feeds into.

We observed significantly lower FRET signal after thermal challenge, even in the case of WT, when comparing the construct with traditional boundaries to one with the N-terminal extension (**Fig 1b and 1c**, scale of Y-axis). Additionally, and strikingly, we observed that Rezatapopt was unable to fully rescue DNA binding of the Y220C truncated construct (**Fig. 1b**). In contrast, inclusion of the 5 additional residues at the N-terminus of the TP53 Y220C DBD conferred full WT-level rescue potential to Rezatapopt (**Fig. 1c**). This parallels observations in more complex biological systems in which full rescue is achievable^5^. These data suggested that this N-terminally extended protein construct was a more biologically relevant model system for studying Y220C behavior.

Further, these results indicate the existence of allosteric communication between residues 89-93 and the Y220C compound binding pocket. The rationale for the allosteric communication observed in **Fig. 1b** and **1c** between residues 89-93 and the small molecule corrector is not immediately obvious. Trp91 is the most N-terminal residue resolved in our structure of the extended Y220C DBD (**Fig 1d**), however, Trp91 and Y220C, where Rezatapopt binds^4^, are on opposite sides of the core β-Sandwich of the DBD (**Fig. 1d**).

Upon closer examination of our Y220C crystal structure of the extended DBD, we observed a similar direct contact between W91 and R174 as was described in the TP53 WT DBD structure previously^12^. This residue is part of the L2 Loop and is directly adjacent to R175, mutation of which is known to be loss-of-function for Zn^2+^-binding^10^. Intriguingly, we observed that R174 is in close enough proximity with H214 for productive contacts to exist. H214 is near the N-terminus of the S7 β-strand leading to the Y220C position. Adjacent to H214 near the N-terminus of the S7 β-strand is R213, which makes a number of intimate N-terminal contacts and coupled with its position on the S7 β-strand, these contacts provide a plausible mechanism for integrating the N-terminal extension of our Y220C DBD construct and the Y220C compound binding pocket. Specifically, the side chain of R213 is positioned for direct contacts with the side chain of S94 and the backbone of S96, residues in close proximity to the N-terminal domain boundary. Further, R213 is also involved in coordinating what appears to be a highly ordered water molecule also present in the boundary-matched WT structure (**PDB 2XWR**). This water molecule appears to structurally integrate the side chain of R213, the backbones of Y163 and M169, as well as the backbone of E171. The latter is particularly of note because the side chain of E171 forms a critical salt bridge with R249^14^. Loss of the E171:R249 salt bridge through the R249S mutation produces an oncogenic variant of TP53, supporting that this intricate network of interactions involving W91, R174, H214, R213, S94, S96, Y163, M169, E171 and R249 is critical for overall protein stability (**Fig 1e**).

Integrating these observations, we hypothesized that there is allosteric communication across the β-Sandwich that integrates the Y220C compound binding pocket, the S7 β-strand, and the N-terminal coil. The focal point of this communication is R213, with W91 being a critical participant (**Fig 1f**). Transmission of conformational information across the domain is compromised in shorter constructs that begin at residue 94, resulting in the divergent behavior of the constructs observed in **Fig. 1b** and **1c**; specifically, the inability of Rezatapopt to fully rescue the truncated TP53 Y220C DBD. The proximity of the intricate structural network in **Fig 1e** to the Zn^2+^ ion immediately poses the question of what role this network plays in the proper organization of the Zn^2+^ coordination site. We therefore set out to interrogate this line of reasoning experimentally and computationally.

### The Oncogenic Y220C Mutation has Compromised Zn^2+^-Binding Affinity

To probe this proposed allosteric connectivity between Y220C and the Zn^2+^, we measured disruption of DNA binding through Zn^2+^-sequestration by EDTA at 25°C. We observed a significant shift in the EDTA IC_50_ on Y220C relative to WT (**Fig. 2a** and **Supplementary Table 1**). To complement this study, we compared the thermal stability of the apo WT and Y220C DBDs. To do this, we utilized EDTA to sequester the Zn^2+^ ion and compared the thermal stabilities of the constructs by DSF. Strikingly, we observed that the change in melting temperature between the WT and Y220C DBDs was eliminated when the Zn^2+^ ion was sequestered by EDTA (**Fig 2a, inset**).

**Figure 2.**
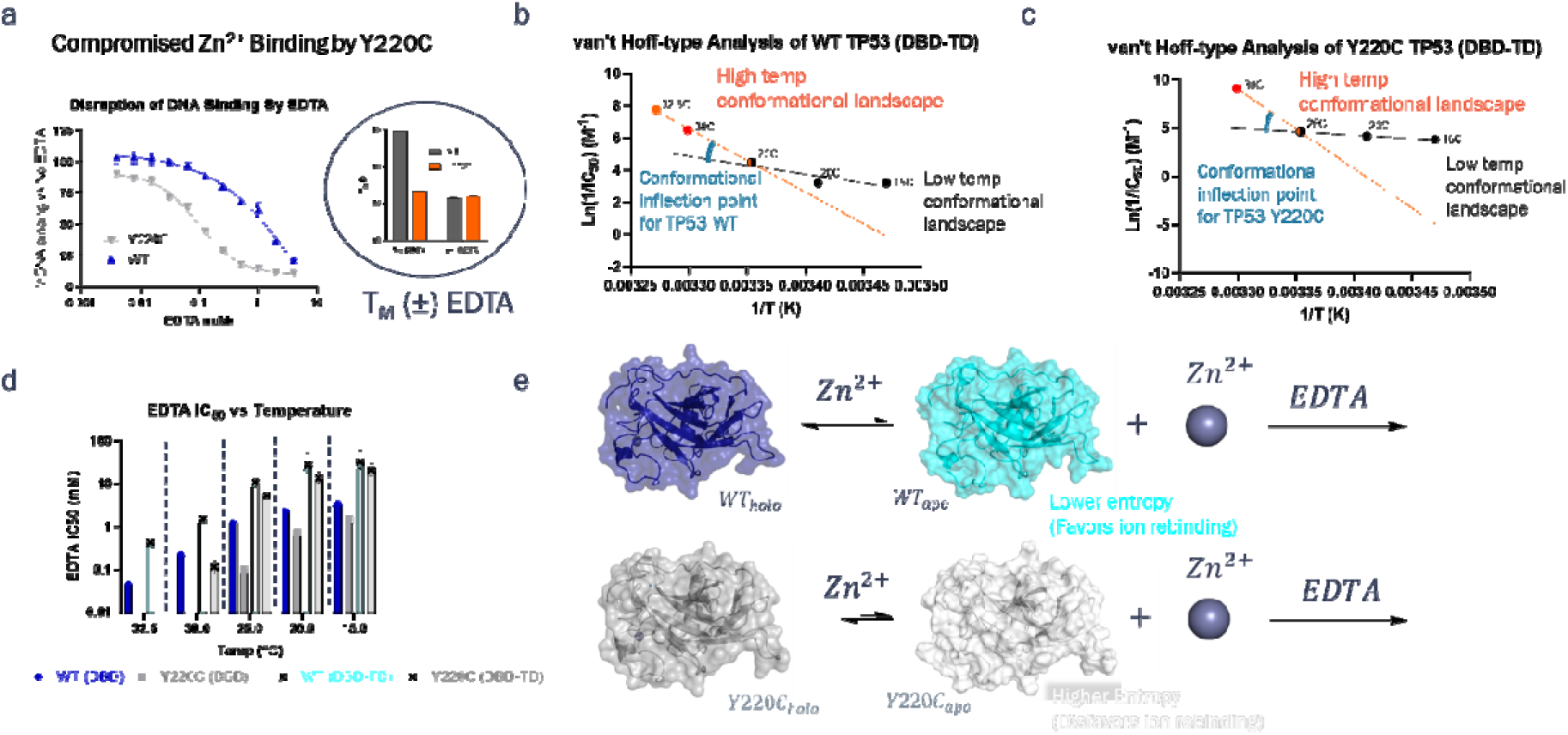
Altered Zinc Affinity between the WT and Y220C DBDs. (**a**) More potent disruption of DNA binding on Y220C DBD vs WT DBD at 25°C; **inset:** Thermal stability comparison by DSF of the Y220C and WT DBDs in the (±) EDTA (WT T_M_: 44.79°C ± .04°C (no EDTA); 35.7°C ± 0.1°C (with EDTA); Y220C T_M_: 36.51°C ± .05°C (no EDTA); 35.94°C ± 0.04°C (with EDTA); duplicate measurements were taken for all conditions). van’t Hoff-type Analysis of (**a**) WT and (**b**) Y220C DBD-TD (DNA binding domain + Tetramerization Domain, residues 89-360). (**d**) Comparison of EDTA IC_50_ values vs Temperature (**e**) Proposed mechanism for altered Zn^2+^-affinity by Y220C DBD relative to WT. **Supplementary Table 1** Provides a summary of all EDTA IC_50_ values.

Surprisingly, these DSF results indicate that the thermal stability difference between the TP53 WT and Y220C DBDs is actually mediated through a loss of Zn^2+^ affinity in the TP53 Y220C DBD. The lack of thermal stabilization of the Y220C DBD by Zn^2+^ and the comparable melting temperature to WT apo DBD indicate that the Y220C mutation does not formally result in thermal destabilization of the DBD per se, but rather, causes loss of affinity for Zn^2+^. In turn, the Y220C DBD is no longer thermally stabilized by Zn^2+^, resulting in a lower observed melting temperature compared to WT.

To thermodynamically probe the destabilization of Zn^2+^-binding by Y220C, we measured disruption of DNA binding through Zn^2+^-sequestration by EDTA at various temperatures. (**Supplementary Figure 2a** and **2b; Supplementary Table 1**). To formally contextualize and interpret this temperature dependence, we utilized a van’t Hoff-type analysis of the EDTA IC_50_ in these experiments as a surrogate for direct assessment of the Zn^2+^-affinity of the TP53 DBD. The concentration range of EDTA used in these experiments is many orders of magnitude above its K_d_ for Zn^2+^ ions, we therefore concluded that changes in the apparent EDTA IC_50_ are not a result of changes in its K_d_ for Zn^2+^ at different temperature, but rather a result of direct changes in the K_d_ of Zn^2+^ with the TP53 DBDs.

The results of this analysis for WT show a linear relationship over the temperature interval of 15°C to 25°C, as expected in a van’t Hoff analysis. However, there is an inflection to a new, linear temperature-dependence from 25°C to 32.5C° (**Supplementary Figure 2c**). Because we are using EDTA IC_50_ values as a surrogate indicator for affinity of Zn^2+^, we cannot make thermodynamic conclusions around changes in the entropy and enthalpy of Zn^2+^ binding to the TP53 DBD. Nonetheless, the inflection in these plots is indicative of a change in the conformational landscape of TP53 that results in increased sampling of dissociation of the structural Zn^2+^, allowing EDTA to sequester the metal ion.

We performed the same analysis on the Y220C DBD but because of the instability of this construct above the key temperature inflection point for WT DBD at 25°C, we could not quantitate the EDTA IC_50_ over the full temperature range. Regardless, we did observe linearity in the van’t Hoff-type plots over the temperature interval of 15°C to 25°C, similar to WT DBD. It is possible the inflection occurs at 20°C for the mutant but is difficult to determine, owing to the limited number of data points (**Supplementary Figure 2c**).

We expanded these studies to constructs containing both the DBD and tetramerization domain to more thoroughly interrogate this potential conformational landscape change (DBD-TD, residues 89-360, **Supplementary Figure 1a**). Our rationale was two-fold: 1) we hypothesized that this larger construct would have additional stability conferred by tetramerization domain-facilitated interactions, thereby allowing us to study the Y220C mutant at temperatures above 25°C; 2) we sought to verify that our observations are likely to be physiologically relevant and not merely an artifact of using the isolated DBD.

We observed an overall increase in stability to EDTA sequestration of Zn^2+^ by both WT and Y220C DBD-TD constructs relative to the respective isolated DBD (**Supplementary Figure 2a, 2b, 2d** and **2e; Supplementary Table 1**). Critically, these larger constructs with more complex equilibria schema in effect from the inclusion of the tetramerization domain still exhibited the same inflection in the WT van’t Hoff plot (**Fig. 2b**). Crucially, we were able to obtain EDTA IC_50_ values at 30°C in this Y220C construct. This allowed us to observe the same evidence for a conformational inflection above 25°C that was seen in the WT DBD construct (**Fig. 2c**). All constructs show evidence of a temperature-dependent decrease in their affinity for Zn^2+^, as evidenced by the improving IC_50_ of EDTA with increasing temperature. Intriguingly, the temperature-dependent difference in EDTA potency shows a wider discrepancy at high temperatures, in contrast to the convergence of IC_50_ values in boundary-matched constructs observed as temperature decreases (**Fig. 2d**)

The convergence of Zn^2+^ affinity between the mutant and WT domains at lower temperatures (**Fig. 2d**) indicates that the holo forms of the WT and Y220C DBDs (*WT_holo_* and *Y220C_holo_*, *respectively*) share a common enthalpic minimum, that is, the stability of the folded, Zn^2+^-bound state is similar. This is supported by the near-perfect alignment of the two domains in their crystal structures and similar HDX shielding patterns (**Supplementary Figure 2f** and **Supplementary Fig. 1d,** respectively).

Therefore, to explain the divergent temperature sensitivity to EDTA of the two domains, we postulated that there are entropic differences between the two domains. Further, because these differences compromise metal binding, we suspected that the entropic differences emerge primarily between the apo states of the domains (WT_apo_ and Y220C_apo_). Entropic differences in this state would be expected to hinder the ability of the metal to associate. Further, this mechanism would allow for the convergence of both domains to a common enthalpic minimum once the metal has bound, thus explaining our EDTA IC_50_ vs. temperature results. In other words, the holo domains collapse to a similar enthalpic minimum observed in the folded crystal structures with divergent responsiveness to EDTA emerging as the energy in the system becomes sufficient for the Zn^2+^-dissociated state to be sampled. Mechanistically, Zn^2+^-dissociation is what permits perturbation of DNA binding by EDTA and behavior of the Zn^2+^-dissociated domains, that is the apo DBDs, would then determine the ability of the metal to rebind the domain rather than be sequestered by EDTA (**Fig 2e**).

### Unbiased & Adaptively Biased Molecular Dynamics Simulations to Illuminate Y220C Loss of Zn^2+^Affinity

To probe this model, we first utilized unbiased molecular dynamics simulations on the apo state of both the WT and Y220C DBDs. Our rationale was that altered conformational dynamics in the Zn^2+^-coordination site would be expected to reduce the ability of the Y220C DBD and Zn^2+^ to productively associate and offer a potential explanation for our experimental results. To begin probing this, we started out with 200 ns trajectories of N-terminally matched structures of both WT (**2XWR**) and Y220C after computationally removing the Zn^2+^ ion and subsequent minimization. The resulting conformational landscapes were then visually inspected for ke potential differences (**Fig. 3a**).

**Figure 3.**
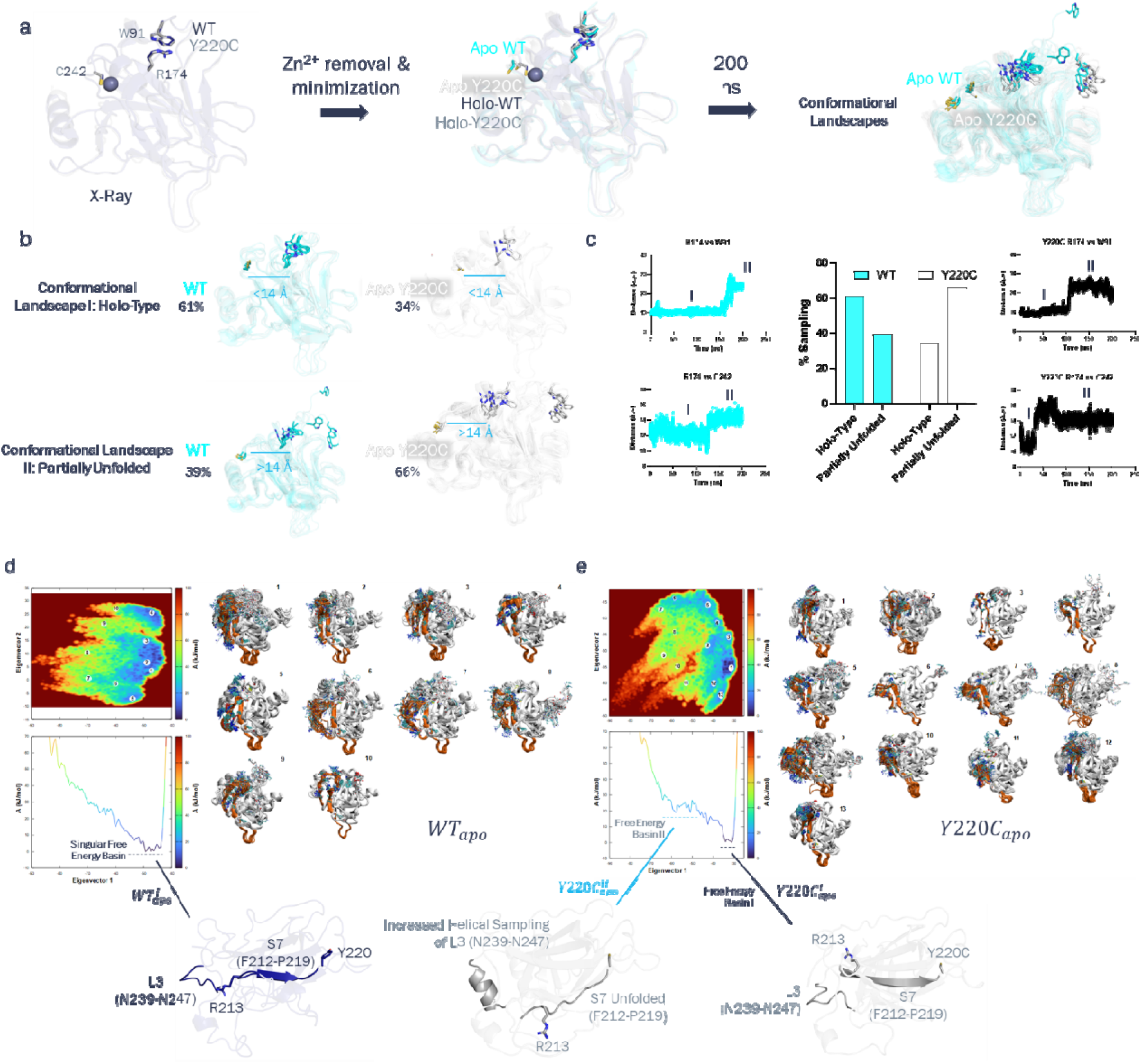
Observation of Altered Conformational Dynamics by the WT and Y220C Apo DBDs in unbiased and Adaptively Biased Molecular Dynamics Simulations. (**a**) Unbiased Molecular Dynamics Workflow. (**b**) Conformational Landscapes sampled by WT and Y220C Apo DBDs. Structures are overlays of the most populated states in each landscape. (**c**) Time evolution of residue spacing between R174 to W91 and R174 to C242 (measured from C_α_) and bar graph depiction of relative time spent in each state. (**c**) and (**d**) Diffusion maps and associated conformational ensembles for apo WT (**c**) and Y220C (**d**) DBDs generated from Adaptively Biased Molecular Dynamics Simulations

Inspection of the Zn^2+^ binding site indicated that C242, which directly contacts the metal, exhibited two distinct populations in both the WT and mutant DBDs (**Fig. 3b**). To quantify this, we classified 2 conformational landscapes defined by the distance between the C_α_’s of C242 and R174. Arg174 was chosen because of its contacts with W91, location in L2, and key role in the interaction network described in **Figure 1e**. The positioning of C242 in **Conformational Landscape I** closely resembles that seen in the Zn^2+^-bound state and is defined by spacing of <14 Å between C242 and R174. **Conformational Landscape II** is marked by an increase in the spacing of C242 and R174 to >14 Å, which significantly distorts the Zn^2+^- coordination site and would therefore be expected to have reduced-to-minimal affinity for the metal ion. Consistent with our implication of W91 in helping to structure the Zn^2+^-coordination site from our experimental results and structural analysis (**Figure 1**), we see substantial displacements of W91 from R174 that are only present in what we have defined as **Conformational Landscape II** (**Fig. 3b**).

Comparison of the time spent in each of these landscapes across the simulations reveals a stark reversal in the relative sampling of these two landscapes by Y220C relative to WT (**Fig 3b** and **3c**). This immediately provides a rationale for the loss of Zn^2+^-affinity by Y220C in which the coordination site spends less time in conformations competent for binding to the metal ion. Plotting of the distances between R174 to either W91 or C242 in both the WT and Y220C DBDs suggests that, not only does Y220C spend more time in **Conformational Landscape II**, the mutant domain also enters into this landscape much more quickly from the initial state. (**Fig. 3c**)

To more thoroughly explore the molecular events linking the metal coordination site with Y220C, we followed up with 2 µsec, Adaptively Biased Molecular Dynamics Simulations^15^. These simulations utilize Adaptive Biasing Potentials to flood the free energy of specific atoms within a molecule in order to better sample the overall conformational landscape and prevent the system from becoming trapped in local minima. By flooding each DBD to the same depth on the same atoms^16^, we can observe differences between the WT and Y220C conformational landscapes, contextualized by the associated free energies of those landscapes.

To reduce the dimensionality of the free energy landscapes, we utilized Diffusion Maps^17^ to estimate the eigenvectors of the Fokker-Planck equation. The simulation trajectories of the biased atoms were then projected for *WT_apo_* and *Y220C_apo_* using these eigenvectors (**Fig. 3d** & **3e**, respectively). Effectively, the Eigenvectors represent a composite of the x,y,z-coordinates of the biased atoms in the simulation. A histogram was constructed for each trajectory along the eigenvector coordinates to estimate the marginalized Boltzmann Distribution, and thus, the free energies: A (kJ/mol).

The key distinguishing feature in the free energy landscapes between the trajectories is a singular free energy basin for WT, contrasted with the sampling of a second free energy basin by Y220C. To distinguish all of the states of the apo domains, we adopted the following nomenclatures: 1) *WT_apo_* and *Y220C_apo_*, in reference to the domains in general, without reference to a specific conformational state; 2) 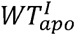, 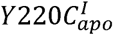 and 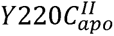 to refer to conformational ensembles within the free energy basins of the respective *WT_apo_* and *Y220C_apo_* DBDs (**Fig. 3d** and **3e**). One of the key distinguishing structural features of the 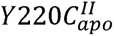 basin is the β-strand character of S7. In this basin, the strand is regularly unfolded. This unfolding has significant influence on R213, which, as discussed earlier (**Figure 1**), appears critical for cooperation with the DBD N-terminus for optimal scaffolding of the Zn^2+^-coordination site. The altered dynamics of this section of amino acids that results from the Y220C mutation provides a direct physical mechanism for the observed allosteric connectivity from the mutated cysteine to the N-terminus and ultimately, the Zn^2+^-coordination site (**Figures 1-3**). Within these simulations, we also observed helical sampling of residues 239-247 of the L3 loop in the 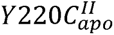 DBD. This segment contains the aforementioned metal-coordinating residue, C242 (**Fig. 3a** and **3b**). Thus, helical sampling of this peptide provides an explicit example of alternate structuring of the Zn^2+^-coordination site uniquely available to *Y220C_apo_*. Further, once helical sampling of the L3 loop is observed, 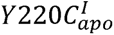 is never revisited in the simulation (**Fig. 3e**). While this does not definitively mean 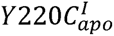 can never be resampled, it indicates the energetic barrier to do so is high. In the *WT_apo_* DBD, the S7 β-strand is much more stably retained throughout the simulation, particularly from residue 215 onward. Further, the helical sampling of L3 observed in Y220C is not observed in the *WT_apo_* simulation (**Fig**. **3d**).

Taken together, these simulations indicate that the Y220C mutation destabilizes the S7 β-strand in *Y220C_apo_*, giving access to distinct conformational states relative to the WT domain in which the Zn^2+^-coordination site of the mutant is altered and unable to bind Zn^2+^. This additional conformational freedom would be expected to result in an overall disfavoring of coordination of the Zn^2+^ ion as proposed in **Fig. 2e**. This is consistent with our experimental results that demonstrate reduced Zn^2+^-affinity as well as our 200ns unbiased simulations of the *WT_apo_* and *Y220C_apo_* DBDs.

### Hydrogen-Deuterium Exchange Studies Support Molecular Dynamics Studies and Indicate Rezatapopt stabilizes Zn^2+^-Binding

In order to experimentally probe the altered conformational landscape of the Y220C DBD, we utilized <u>H</u>ydrogen-<u>D</u>euterium Exchange (HDX). In short, we monitored deuterium uptake after temperature challenge for 1, 30 and 60 minutes in TP53 Y220C DBD bound to Rezatapopt relative to a DMSO control. We performed these studies at 30°C, above the observed critical inflection in our temperature-dependent biochemical studies, but crucially below the measured T_m_ of the Y220C domain. Our rationale for these studies was that shielding effects by the compound over time could map out the sequence of conformational changes occurring in the DMSO-treated Y220C DBD (**Fig. 4a**).

**Figure 4.**
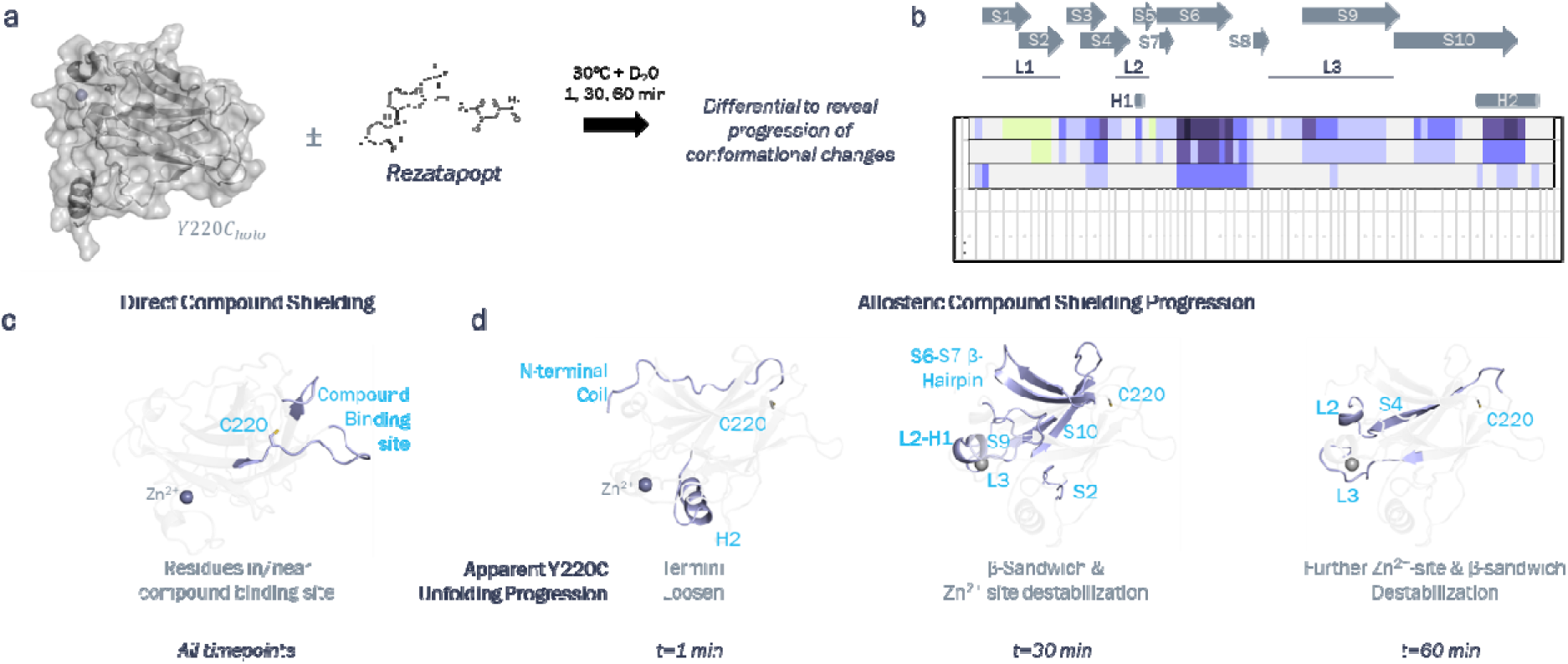
HDX Differential analysis of TP53 Y220C DBD Unfolding (±) Rezatapopt. (**a**) Experimental set up between DMSO-treated TP53 Y220C DBD and Rezatapopt-treated after 1, 30 and 60 min of challenge at 30°C followed by cooling to RT and 10 sec labeling in D_2_O. (**b**) Differential shielding between DMSO-treated TP53 Y220C DBD and Rezatapopt-treated. Secondary structural elements contained within respective peptides are shown (**c**) Direct compound shielding effects mapped onto TP53 Y220 DBD coordinates shown in **Supplementary Fig. 1f**. (**d**) Evolution of new allosteric shielding events observed at each time point mapped onto TP53 Y220 DBD coordinates shown in **Supplementary Fig. 1f**.

Analysis of the shielding differentials (**Fig. 4b**) over time yielded an intriguing mix of both direct and allosteric effects of the compound. At the earliest time point, we observed predominantly shielding in peptides surrounding the compound binding site. These shielding events only deepened throughout the course of the experiment. Unfortunately, we are unable to distinguish whether these peptides are exhibiting altered mobility and deuterium uptake or whether the shielding is entirely due to steric occlusion from being in direct contact with the compound (**Fig. 4c**).

We observed allosteric shielding in the N-terminal residues at 1 min (**Fig. 4d**), biophysically demonstrating the allosteric communication between residues 89-93 observed in our studies in **Figure 1**. This shielding was not robustly maintained across all time points (**Fig. 4b**), indicating this region has its conformational mobility reduced, but not fully restricted by the compound. We also observed robust shielding in H2, which is part of the DNA-contact surface stabilized by Zn^2+^. Taken with our EDTA data in **Figure 2** and **Supplementary Figure 2**, this may be the first sign of the reduced Zn^2+^-affinity where the metal ion has dissociated from the protein, allowing more conformational freedom to H2 (**Fig 4d**, **t=1 min panel**).

At the 30-minute time point, we see increased deuterium uptake in S2 that we believe to be related to its proximity to H2, which showed increased flexibility at the first time point. We also observed a particularly complex cascade of allosteric changes involving both the central β-sandwich and the Zn^2+^-coordination site, with stabilization by the compound in the S6-S7 β-Hairpin, as well as in S9 and S10. In the Zn^2+^-coordination site, we observed compound shielding emerging in the L2 and L3 loops as well as H1 (**Fig. 4d**, **t=30 min panel**).

At the final time point, we see new shielding events in peptides in the L2 and L3 loops as well as the centrally located S4 (**Fig 4d**, **t=60 min panel**). The broad increased uptake of the DMSO-treated Y220C DBD is consistent with it existing in a conformational landscape similar to the Y220C^II^ landscape seen in the molecular dynamics simulations. While we do not have the temporal resolution in these studies to fully delineate the sequence of events contributing to the shielding at each time point, these studies are consistent with our Adaptively Biased Molecular Dynamics simulations where S4 is stable throughout the WT simulation and there is a correlated destabilization of the structure of especially the S6-S7 β-Hairpin containing R213 and the Zn^2+^-coordination site in Y220C (**Fig. 1d-f** and **Fig 3e**).

Intriguingly, these studies directly demonstrate allosteric stabilization of the Zn^2+^-binding site by Rezatapopt (**Fig. 4d**), suggesting that Rezatapopt functions to restore the Zn^2+^ binding affinity loss bestowed by the Y220C mutation. We therefore set out to understand the thermodynamic binding mechanism of Rezatapopt to Y220C with respect to Zn^2+^.

### Rezatapopt Restores WT-Level Zn^2+^ affinity to Y220C DBD

To probe the binding mechanism of Rezatapopt with the Y220C DBD and Zn^2+^, we first looked at its ability to preserve Y220C DNA binding in the presence of EDTA at 20°C. Consistent with the HDX studies showing stabilization of the Zn^2+^-site by Rezatapopt, we observed that Rezatapopt weakens the EDTA IC_50_ for disruption of Y220C DNA binding in a dose-dependent fashion (**Fig. 5a**; **Supplementary Table 2**).

**Figure 5.**
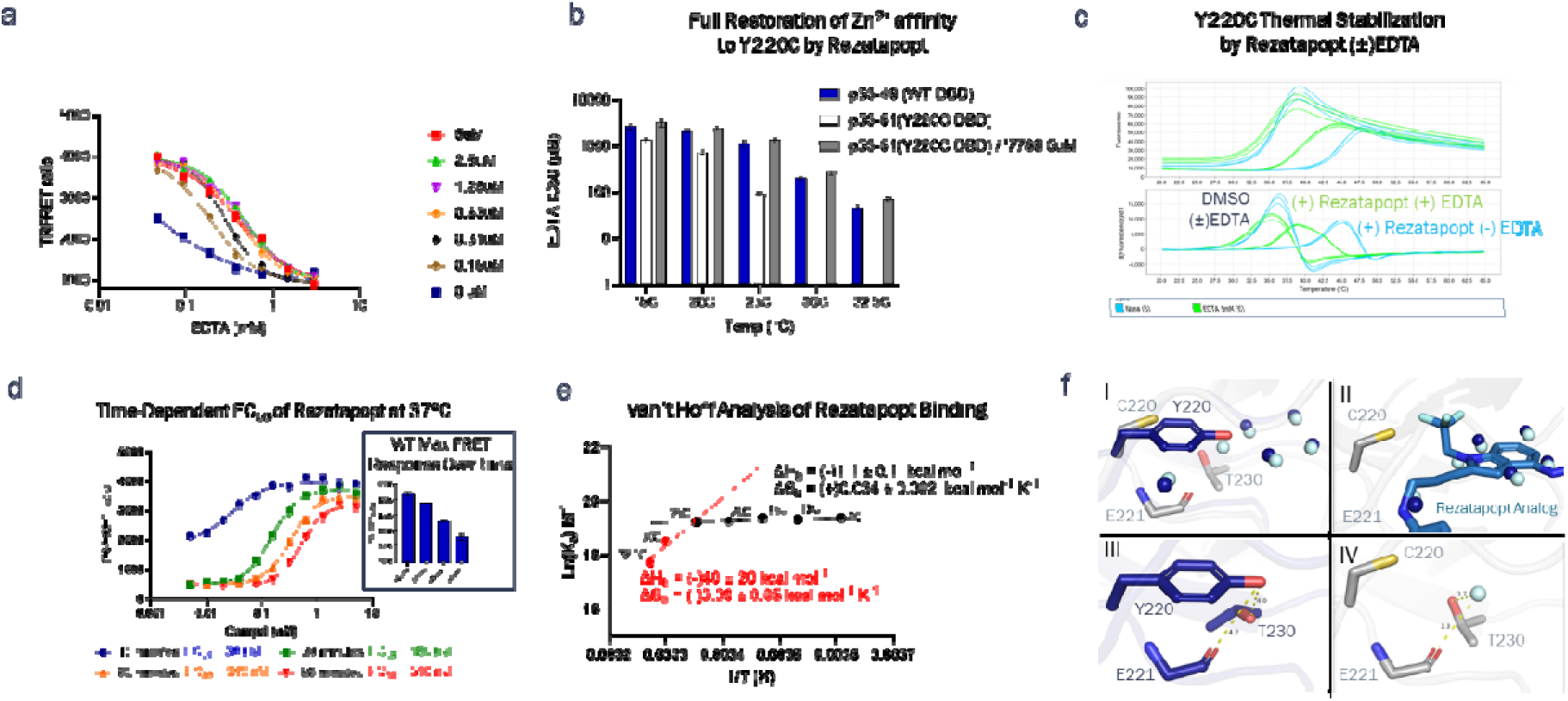
Binding Mechanism Studies with Rezatapopt. (**a**) Dose-Dependent Rescue of EDTA-sequestration from Y220C DBD by Rezatapopt (Values and error summarized in **Supplementary Table 2**). (**b**) IC_50_ of EDTA for disruption of DNA binding at various temperatures and restoration of WT-level of Zn^2+^ affinity to TP53 Y220C DBD by Rezatapopt (values and error summarized in **Supplementary Table 3**). (**c**) Reductions in thermal stabilization of the Y220C DBD by Rezatapopt as a result of Zn^2+^ sequestration by EDTA. (DMSO T_M_: 36.14°C ± 0.09°C; DMSO + EDTA T_M_: 35.7°C ± 0.2°C; Rezatapopt T_M_: 45.68°C ± 0.04°C; Rezatapopt + EDTA T_M_: 38.5°C ± 0.1°C; all measurements were taken in duplicate) (**d**) Time-Dependent Shift in Rezatapopt rescue of Y220C EC_50_ at 37°C. Samples were initially incubated together at RT and the shift in EC_50_ reflects the change in overall equilibrium as the samples equilibrate to 37°C (values and error summarized in **Supplementary Table 4)** (**e**) van’t Hoff Analysis of Rezatapopt Binding to TP53 Y220C DBD. Δ_b_H=(-)1.1 ± 0.1 kcal mol^−1^K^−1^, Δ_b_S= (+)0.034 ± .002 kcal mol^−1^K^−1^ over the low temp interval and Δ_b_H=(-)40 ± 15 kcal mol^−1^K^−1^, ΔS_b_= (-)0.08 ± 0.05 kcal mol^−1^K^−1^ over the high temp interval (values and error summarized in **Supplementary Table 5**). (**f**) Structural evaluation of potential ordered water molecules near residue 220 in WT and mutant DBDs (Dark blue = WT, Gray=Y220C, Y220C water molecules = cyan, WT water molecules = Dark blue).

To compare the Zn^2+^ affinity of Rezatapopt-bound Y220C relative to the WT DBD, we evaluated the EDTA IC_50_ for disruption of DNA binding across the same temperature range as our previous experiments described in **Figure 2**. We utilized 5 µM Rezatapopt because this was sufficient to maximize Zn^2+^-stabilization to EDTA (**Fig. 5a**; **Supplementary Table 2**). We observed congruence with respect to EDTA IC_50_ between the WT and Rezatapopt-bound Y220C DBDs, indicating that Rezatapopt-bound Y220C has the same potency for Zn^2+^ as the WT DBD (**Fig. 5b**; **Supplementary Figure 3** and **Supplementary Table 3**).

Our experimental results described in **Figures 1, 2** and **5** combined with the MD studies in **Figure 3** strongly argue that the Y220C DBD is severely, if not irreparably, impaired for binding Zn^2+^ at physiological temperatures. Thus, we hypothesized that the cellular mechanism of action of Rezatapopt may be to bind and stabilize *Y220C_apo_* and facilitate ion loading. Therefore, we undertook a series of studies to interrogate this hypothesis.

First, we probed for the ability of Rezatapopt to bind *Y220C_apo_* using DSF in the presence of EDTA. We observed stabilization of *Y220C_apo_* bound to Rezatapopt in the presence of EDTA, supporting our hypothesis that the compound can engage the apo domain (**Fig. 5c**). Further, the reduced T_m_ is suggestive that the affinity is reduced for apo state of the domain as would be expected, given the cooperative nature of the compound and the metal ion (**Fig. 5a**). Thus, one would expect a drastically different potency for the interaction between Rezatapopt and Y220C at temperatures where the Zn^2+^ is destabilized.

To probe this, we first turned to our DBD-TD construct to enable us to study Rezatapopt potency at 37°C. We prebound Rezatapopt, the Y220C DBD-TD, and DNA at room temperature to allow the complex to stably form. We then incubated the samples at 37°C and monitored the Rezatapopt potency at varying time points. We observed a significant weakening of the Rezatapopt EC_50_ over time. We attribute this to the establishment of new equilibria involving Zn^2+^ dissociation and Rezatapopt binding to the apo DBD as the temperature in the samples increased (**Fig. 5d; Supplementary Table 4**). Of note, the max FRET response for both the WT and Y220C-Rezatapopt samples decreased over time (**Fig. 5d+inset**). Our interpretation of this behavior relates to the *WT_apo_* T*_m_* of ∼36°C (**Fig. 2a**): with each cycle of Zinc unbounding and rebinding, ∼30-50% of the sampled *WT_apo_* population spontaneously samples an unfolded state and some fraction is lost to irreversible denaturation (discussed and visually depicted later).

To more rigorously probe the altered affinity of Rezatapopt at higher temperatures, we utilized a TR-FRET based probe displacement assay using a biotinylated Rezatapopt derivative to directly assess compound binding affinity at different temperatures (**Supplementary Figure 4a**). We restricted these studies to the DBD. This was intentional in order to avoid any confounding influences from tetramerization. Further, in contrast to our previous temperature dependent studies, it is worth noting that there is no EDTA nor DNA in these compound experiments (**Figure 2**), ruling out more complex behavior involving confounding equilibria from these species in our previous experiments. To enable these studies, the probe EC_50_ was determined at a range of temperatures to estimate its K_d_. Competition assays were run with the probe concentration normalized to its observed EC_50_/K_d_ at that respective temperature (**Supplementary Figure 4b**). Rezatapopt IC_50_ values at each temperature were determined using this probe displacement assay and IC_50_ values were converted to K_d_ through the Cheng-Prusof relationship^18^. To validate this protocol, the K_d_ of Rezatapopt at 20°C was confirmed by Surface Plasmon Resonance (SPR) with precise agreement between this biophysical technique and the probe displacement assay at 20°C, validating this procedure for K_d_ determinations (**Supplementary Figure 4c-f**). The full K_d_ vs temperature relationship was not accessible through SPR due to slow binding kinetics at low temperatures preventing full equilibration of binding. Additionally, Y220C DBD surface instability was observed at higher temperatures, further disqualifying this technique for this analysis.

We found compound potency changes were minimal over the range of 4°C to 25°C. At temperatures above 25°C, there is a sharp deflection downward in the van’t Hoff plot, reciprocal to what was seen with EDTA-mediated disruption of DNA binding (**Fig 5e**; **Supplementary Figure 5** and **Supplementary Table 5**). Of note, because the studies in **Figure 5e** constitute a proper van’t Hoff analysis in comparison to the studies in **Figure 2**, conclusions can be drawn about both the entropy and enthalpy of binding. Accordingly, in the lower temperature regime, binding is minimally exothermic with a positive entropic contribution to binding (ΔH_b_ = −1.1 ± 0.1 kcal mol^−1^; ΔS_b_ = 0.034 ± 0.002 kcal mol^−1^K^−1^; **Supplementary Table 5**). One rationalization of these values is that binding is a function largely of the hydrophobic effect with both the binding pocket and small molecule being highly hydrophobic. This could lead to an increase in entropy upon binding as a result of release of highly ordered water molecules in the binding pocket and around the small molecule.

Intrigued by this, we examined the conserved water network around residue 220 in the WT and Y220C structures (**2XWR** and XXXX, respectively). We overlaid the structure of a Rezatapopt analog we had determined in our Y220C crystallography system that shows analogous, entropically-driven binding at low temperatures (**Supplementary Figure 5**, Compound 3; **Supplementary Table 5**). All of the water molecules shown in the network were displaced by the Rezatapopt analog (**Fig. 5f, panels I & II**), providing a plausible structural explanation for the entropically-driven binding of Rezatapopt and its analogs at low temperatures.

Further, in the non-compound bound state, we identified one water molecule that was unique to the Y220C mutant in the immediate vicinity of residue 220. Likely not coincidentally, this water is positioned very similarly to the -OH in the WT Y220 residue and is in close enough proximity to both the sidechain of T230 and the backbone of E221 to form hydrogen bonds. It is possible this water is a key structural water in the mutant by acting as a surrogate for the hydroxyl of Y220 at low temperatures and liberation/destabilization of this water at increasing temperatures is a key contributor to the exaggerated conformational inflection in the Y220C DBD relative to the WT DBD. (**Fig. 5f, panels I, III** and **IV**).

With respect to Rezatapopt binding (**Fig. 5e; Supplementary Table 5**), at temperatures above the inflection point in the van’t Hoff plot, binding becomes entirely enthalpically driven with the ΔS_b_ estimated to be negative (ΔS_b_ = (-)0.08 ± 0.05 kcal mol^−1^K^−1^). We rationalize this as follows: the switch from positive to negative entropy is consistent with a transition to a mechanism where binding of the Zn^2+^ ion is restored, resulting in a reduction in the overall entropy due to the restoration of complexation. Further, binding of the Zn^2+^ would explain the exceptionally large ΔH_b_ estimate in this temperature range [(-)40 ± 20 kcal mol^−1^], which is difficult to rationalize for the binding of Rezatapopt to Y220C, particularly when the measured enthalpy of binding is so minimal at lower temperatures. The shift to highly favorable enthalpy presumably requires new contacts to be made with the domain, which is unlikely for the compound, at least to the extent required to explain the difference in enthalpic contributions between the two temperature ranges, particularly given that binding is weakened, not strengthened, at higher temperatures. However, this is rationalizable if the enthalpic contacts are coming from the Zn^2+^ ion’s restored contacts with its coordination site.

Finally, binding of the compound to the *Y220C_apo_* DBD that restores Zn^2+^ affinity would be expected to reduce the conformational entropy of *Y220C_apo_*, providing an additional/alternative rationale for the transition to a negative ΔS_b_ (**Fig. 5e**). We performed the same analysis on two Rezatapopt analogs, including the one mentioned previously for structural studies. These 3 analogs cover an ∼50-fold range of affinities. In all 3 instances, we observed analogous behavior in the van’t Hoff plots, indicating this behavior is a general phenomenon of this chemotype and likely, to most, if not all, Y220C reversible correctors (**Supplementary Figure 5** and **Supplementary Table 5**).

To summarize the scope of our learnings throughout the course of these studies, we have proposed a series of free energy diagrams to offer holistic interpretations of our datasets and the relevant macroscopic states of the WT and Y220C p53 DBDs spanning from physiological temperature to temperatures in the typical experimental range. Positioning along the vertical axis (ΔG) in these diagrams is intended to convey relative energy differences and should not be interpreted as precise quantitative assessments of the relative free energies. We have layered onto these coordinates specific experimental and computational data points from the preceding discussion to help contextualize each state in the coordinate.

We propose three different contributing ensembles for WT Conformational Free Energy Landscape in relation to Zn^2+^ Binding (**Fig. 6a**). There are two thermodynamic sinks within the diagram. The left most sink is full denaturation of the DBD 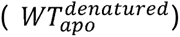 that siphons the protein over time (**Fig. 5d, inset**). This is only expected to be meaningfully observable at temperatures >25°C (Conformational Inflection Point). At temperatures above the inflection point, a significant fraction of the WT DBD exists in an equilibrium of folded 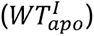 and unfolded 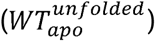 states (T*_m_* ∼36°C). Our Adaptively Biased MD studies indicate the 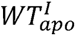 exhibits a conformational ensemble with a singular free energy basin and in conjunction with our experimental data, this ensemble is capable of metal ion coordination. Metal complexation dramatically thermally stabilizes the domain (T_m_ 45°C, 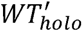). This interplay explains our observed potent DNA binding with the slow loss of signal over time at 37°C (**Fig. 5d inset**), as well as the steep increase in EDTA IC_50_ for zinc sequestration above the inflection point. Below the inflection point, the system is dominated by the metal complexed DBD with an ordered water network around Y220 stabilizing that region of the protein and limiting conformational mobility (*WT_holo_*). At ∼25°C (inflection point in van’t Hoff plots), we propose this water network becomes destabilized, which would increase the conformational mobility around Y220 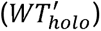. Our MD studies indicate that increased conformational flexibility in this region destabilizes S7, which destabilizes R213, ultimately destabilizing the Zn^2+^-coordination site. By contrast, at temperatures below the inflection point, the water network is more stably maintained. Thus, the domain populates the second thermodynamic sink and the probable enthalpic minimum (*WT_holo_*). The stable water network limits the conformational sampling of the DBD at low temperatures, thus tightly securing the Zn^2+^ in its coordination site, experimentally observable through the less potent EDTA IC_50_ at low temperatures. Importantly, at all temperatures, one would expect each of these states to contribute to the macroscopic behavior of the protein with the relative populations of each being modulated by the temperature of the system (**Fig. 6a**).

**Figure 6.**
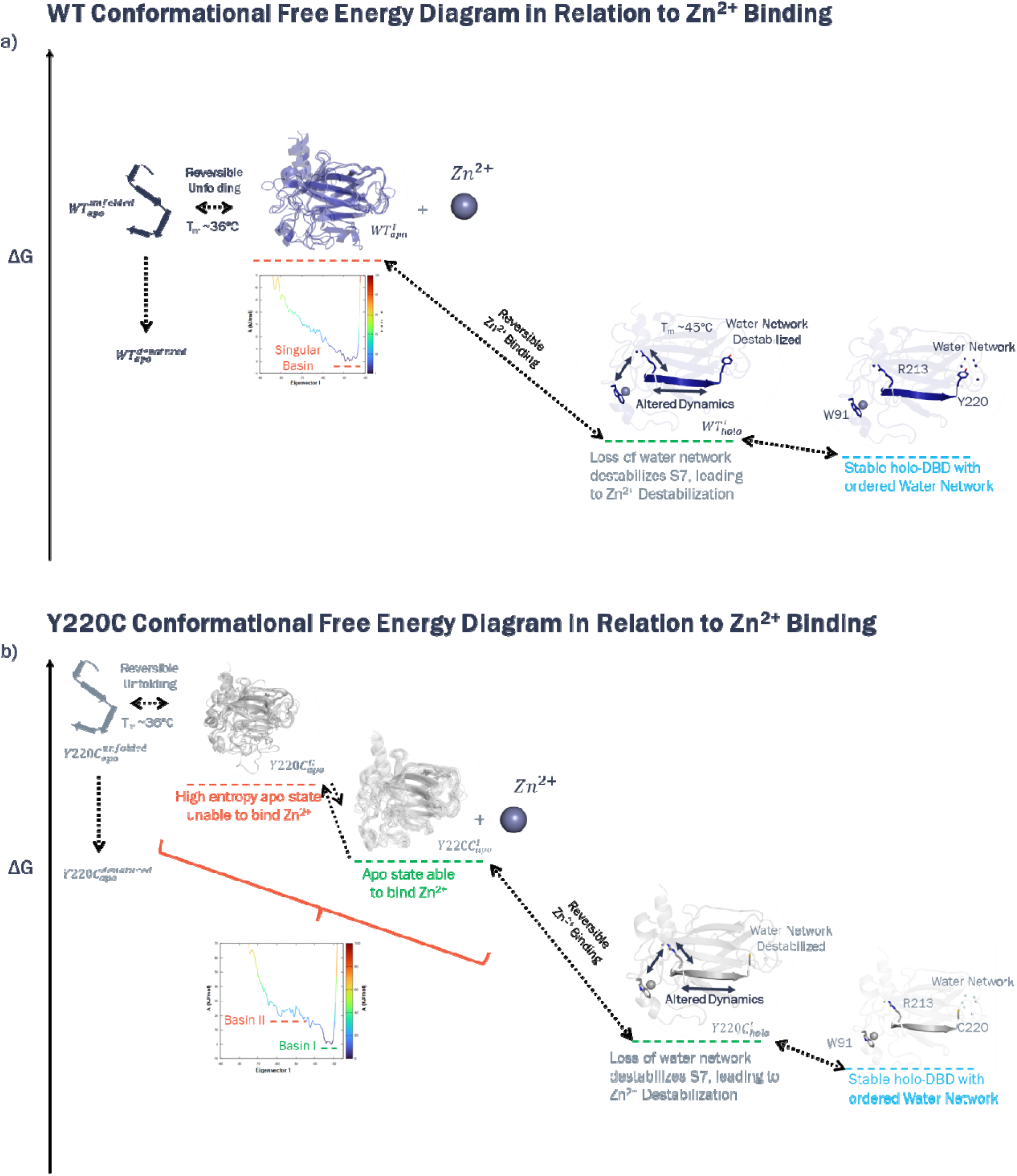
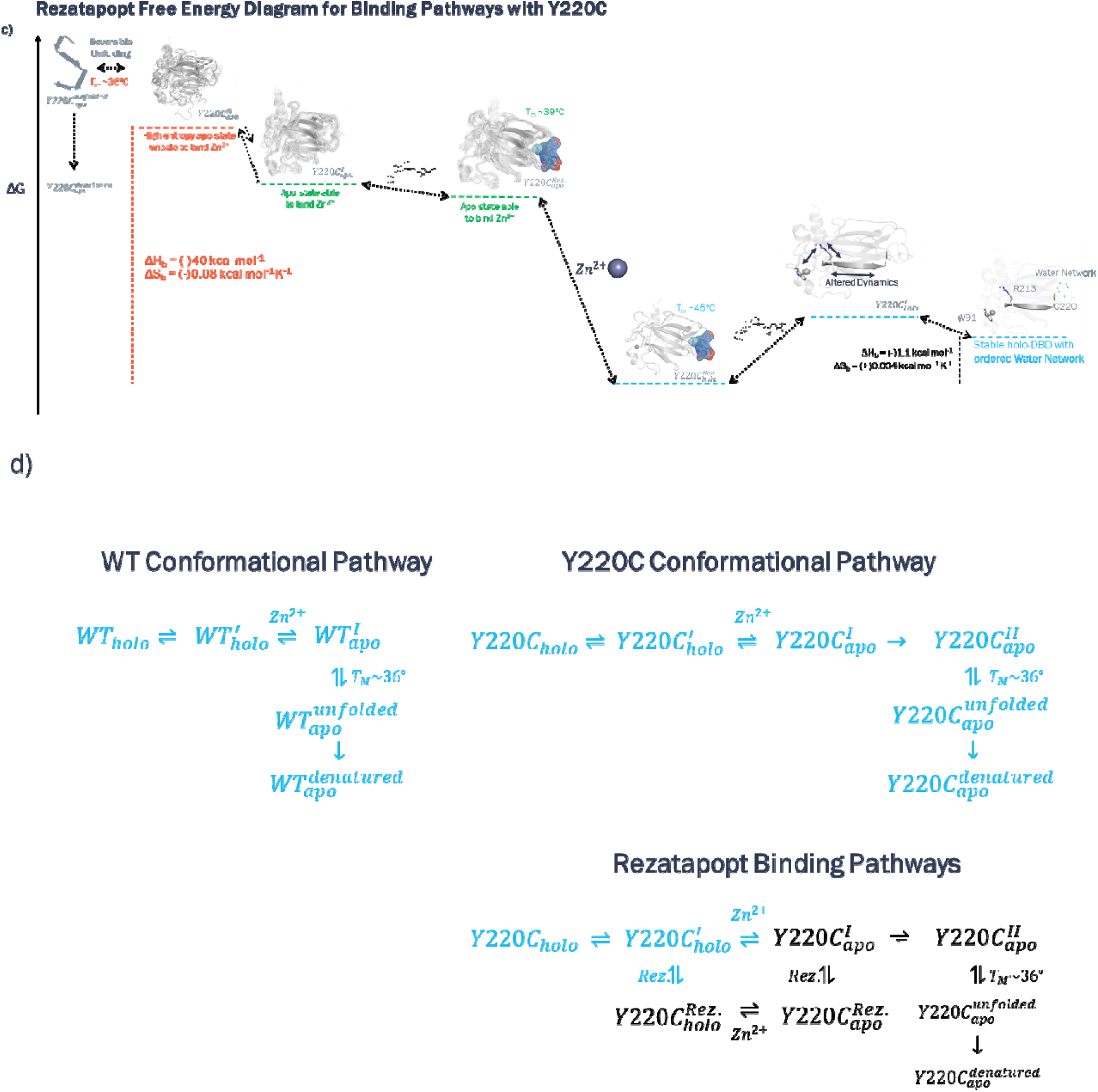
Integrated Mechanistic Interpretations of the WT and Y220C Conformational Pathways and the Rezatapopt Binding Mechanisms. Proposed Relative Reaction Coordinates for the Conformational Equilibria of the WT (**a**) and Y220C (**b**) DBDs and for the Multiple Rezatapopt Binding Pathways (**c**). Relative free energy differences are qualitatively depicted and not meant to convey absolute energy differences. Crystallographic, DSF and Molecular Dynamics Observations are integrated with the temperature dependence of the (DBD-TD) Zn^2+^ affinities measured from Figure 3a and **3b** to illustrate the relevant conformational ensembles believed to contribute to the van’t Hoff plots. (**d**) Proposed thermodynamic mechanisms for the relevant pathways summarizing the free energy coordinates depicted in (**a-c**). Shown in black is the expected mechanistic pathway relevant at physiological temperature.

The Y220C Conformational Free Energy Landscape in Relation to Zn^2+^ Binding is very similar to the WT with the following key differences (**Fig. 6b**): 1) At physiological temperatures, a second conformational free energy basin is accessible to the domain owing to increased conformational freedom from the smaller C220 side chain that destabilizes S7, giving rise to an additional conformational state/ensemble that is incapable of binding Zn^2+^ (**Fig. 2a**, 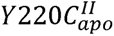). The totality of our data indicate this is the near exclusive conformational ensemble of the protein at physiological temperatures; 2) The available conformational pathway to 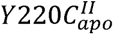 exaggerates the Zn^2+^ affinity difference between WT and Y220C at temperatures approaching, equal to and greater than the inflection point; 3) The Y220C DBD water network in the enthalpic minimum, *Y220C_holo_*, contains an additional water as a surrogate for the tyrosine -OH (**Fig. 5f**). The displaceable nature of this molecule compared to the amino acid side chain contributes substantially, if not entirely, to exaggerating the Zn^2+^ affinity differences between the mutant and WT domains as temperatures approach the inflection point. When temperature is sufficiently low, the water network is stabilized in both the Y220C and WT domains such that they approach a common enthalpic minimum and display nearly identical affinities for Zinc (**Fig. 6a** and **6b, Supplementary Table 1**).

The 3^rd^ diagram we propose is with respect to the Rezatapopt Free Energy Coordinate for its available binding pathways with Y220C (**Fig. 6c**). Rezatapopt is likely able to ingress into the Y220C Conformational Coordinate at either 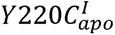 or 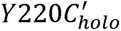. At physiological temperatures, the state most available to it is 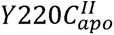, however, overlay of the structure with a Rezatapopt analog with states in 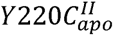 reveal expected clashes that would preclude compound binding (**Supplementary Figure 6**). We therefore propose a conformational selection mechanism in which Rezatapopt binds to the limited pool of sampled 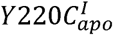 at physiological temperatures and shifts the overall equilibrium through mass action. Binding to this state collapses the domain back to a WT-like apo state and gives a thermodynamic signature of a T_m_ of ∼39°C (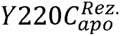, lower in energy than the compound-free apo state, 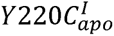, **Fig. 6b** and **6c**). Populating 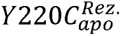 restores metal binding, resulting in a large enthalpy of binding from metal coordination and negative entropy from the formation of the ternary 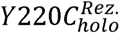 (**Fig. 5e**). Another pathway is also available from this state in which Rezatapopt can dissociate and both 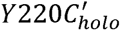 and *Y220C_holo_* are sampled. Above the inflection point, this is not a significant contributor because the instability of the water network destabilizes these states. Below the inflection point however, this pathway becomes dominant. The dissociation of the molecule allows the conserved water network to stably form. At low temperatures, the forward direction of this pathway for binding of Rezatapopt is favored entropically, driven through liberation of the ordered waters, despite the minimal enthalpic contributions observed in the thermodynamic signature of the binding event. While both pathways are formally expected to be operative in any temperature range, the overwhelming differences in the stabilities of the Y220C DBD states above and below the inflection point dictate that only one pathway effectively contributes to the binding at a given temperature with the exception of temperatures near the inflection point, where both pathways likely contribute similarly, and thus, explaining the mechanistic shift in the van’t Hoff plot.

Finally, we condensed each of these into the thermodynamic mechanisms show in **Figure 6d**.

## Conclusions

Our studies here represent, to our knowledge, the first systematic recombinant exploration of the Y220C DBD utilizing both the endogenous sequence and the more biologically relevant domain boundaries. These studies robustly demonstrate a temperature-dependent compromised ability of the Y220C DBD to bind Zn^2+^ relative to the WT DBDs.

While our initial intent was simply to develop a more robust assay system for biochemical studies by increasing protein stability through the N-terminal extension of the DBD, doing so led to the series of biochemical, biophysical, structural and molecular dynamics studies described here to reveal detailed insights into the conformational landscape of both the Y220C and WT DBDs. These insights involve the illumination of residue-level detail into previously unrecognized allosteric connectivity across the TP53 DBD. This connectivity is temperature modulated, which appears to be mediated through the stability of an ordered water network present around residue 220. The Y220C mutation creates a void in the protein that appears to be filled by a water molecule that substitutes for the -OH of the tyrosine side chain in the WT. This displaceable water molecule exaggerates the ability of temperature to modulate the allosteric connectivity across the domain, resulting in an exacerbation of the temperature-dependent decrease in metal-affinity of the WT DBD. Once dissociated from the metal, the smaller side chain of the Y220C DBD allows for sampling of additional conformational states that reduces the likelihood of metal binding, leading to the overall instability of the protein at physiological temperatures.

Rezatapopt is able to correct the conformational ensemble of the Y220C DBD to restore WT-level affinity for the metal at all temperatures tested. The high enthalpic contribution of Rezatapopt binding at high temperatures is best explained through the restoration of metal complexation, which could also explain the negative entropy of binding. Further, folding of the domain as a result of restored metal complexation would further contribute to a negative ΔS_b_. This is in contrast to the positive entropy and limited enthalpic contributions observed at low temperatures where metal binding is stabilized through displacement of the conserved water network.

Accordingly, one of the key implications of this work is that an optimization campaign focused on our proposed key physiological state of Y220C, namely 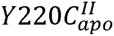, could lead to chemical matter with improved cellular potencies. The apo state of Y220C has been seldom considered in the design of small molecule correctors and our work suggests that targeting of the Y220C apo state of the domain could allow access to forms of TP53 with superior stability compared to the WT protein. Specifically, increasing of the thermal stability of the apo domain could limit the gradual loss of protein observed in our studies through unfolding and denaturation at physiological temperatures. Alternatively, enhancement of Zn^2+^ affinity beyond that of WT could also be used to pursue a pharmcologically-induced, “superstable” DBD.

Finally, while the small molecule binding potential is limited around the Y220 region in WT, our results do suggest that intervention at this region could be used to tune the affinity of the Zn^2+^ ion, potentially opening up targeting of other TP53 missense mutations.

**Accession IDs:**

**TP53:** P04637

## Experimental Procedures

### Protein expression and purification

TP53 vectors (**Table 1**, Uniprot ID: P04637) with His_6_- or His_6_- and Flag- or Avi- tags were transformed into competent BL21 (DE3) gold cells. Cells were grown in LB medium supplemented with 100 µM ZnCl_2_ to an OD 0.6-0.8 and protein expression was induced with 500 μM isopropyl β-D-1-thiogalactoyranoside (IPTG) at 16°C for 20 h. Cells were harvested by centrifugation (4000 x g, 30 min, 4°C), bacteria were lysed by sonication (400 W, 3 sec pulse on, 3 sec pulse off for 30 min on ice) in 50 mM Tris, pH 7.2, 5 mM DTT, 150 mM NaCl, 5% glycerol, 1 mM PMSF (10ml/g cell paste). Lysates were cleared by centrifugation (20,000 x g, 30 min, 4°C), the supernatant was collected and cleared for a second time. The supernatant was applied onto pre-equilibrated Ni-NTA beads (Roche, 3 mL/1 L cell culture), washed with lysis buffer and lysis buffer with 20 mM imidazole before eluting the target protein with lysis buffer containing 250 mM imidazole. His-tags were cleaved with TEV protease in a 40:1 ratio of protein: protease during dialysis against lysis buffer over night at 4°C and removed with another round of Ni-NTA purification collecting the target protein in the flow-through. For TP53(89-293, Y220C) crystallography, the sample was subsequently passed over a Heparin column (GE, Buffer A: 50 mM Tris pH 7.2, 25 mM NaCl, 5 mM DTT, Buffer B: 50 mM Tris pH 7.2, 1 M NaCl, 5 mM DTT) and finally over size-exclusion (GE, S75, 24 mL), before being flash frozen in SEC/storage buffer (25 mM KH_2_PO_4_, pH 7.2, 150 mM NaCl, 5 mM DTT, 5% Glycerol). Protein molecular weight was confirmed for each construct with intact LC/MS.

**Table 1.** TP53 protein constructs used in this study.

| Vector | N-term tag | Boundaries | Mutations | C-term tag |
| --- | --- | --- | --- | --- |
| pET28a | His<sub>6</sub>-TEV | 89-293 | Y220C | N/A |
| pET28a | His<sub>6</sub>-TEV | 89-312 | N/A | GGSGGS-FLAG |
| pET28a | His<sub>6</sub>-TEV | 89-312 | Y220C | GGSGGS-FLAG |
| pET24a | His<sub>6</sub>-TEV | 94-312 | N/A | GGSGGS-FLAG |
| pET24a | His<sub>6</sub>-TEV | 94-312 | Y220C | GGSGGS-FLAG |
| pET24a | His<sub>6</sub>-TEV | 94-312 | Y220C | GGSGGS-AVI |
| pET28a | His<sub>6</sub>-TEV | 89-360 | N/A | GGSGGS-FLAG |
| pET28a | His<sub>6</sub>-TEV | 89-360 | Y220C | GGSGGS-FLAG |

### Crystallization and structure determination of TP53(89-293, Y220C)

His-tag cleaved TP53 (G-P89-G293, Y220C; 4 mg/mL or ∼43 μM) was kept as-isolated or incubated with **Compound 3** (1 mM) for 2 hr (4°C) before crystallization. Data-quality crystals were grown using the sitting-drop vapor diffusion method at room temperature by mixing 200 nL of protein mixture with 200 nL of precipitant over a reservoir of 200 μL of precipitant (50 mM ammonium sulfate, 15% (w/v) PEG 8000, 100 mM sodium citrate (apo) or 200 mM ammonium sulfate, 100 mM BIS-TRIS pH 6.5, 25% (w/v) PEG 3350 (**Compound 3**)). Crystals were cryoprotected with crystallization precipitant supplemented with 20% (v/v) glycerol before being flash frozen in liquid nitrogen prior to data collection. X-ray diffraction data were collected at the Shanghai Synchrotron Radiation Facility (Pudong, China) on beamline BL19U1 and processed in the space group P2_1_ using XDS^19^. All crystallographic refinement tools were accessed through the CCP4 suite^20^. The initial structure of TP53 was determined by molecular replacement in PHASER using the protein coordinates of PDB ID 2XWR. Ligand restraints were generated using JLigand and AceDRG and structure refinement was carried out in Refmac5 combined with several rounds of manual fitting in COOT. Figures were prepared with PyMOL (The PyMOL Molecular Graphics System, Version 3.0 Schrödinger, LLC). Complete data collection and refinement statistics are shown in **Supplementary Table 6** and **Supplementary Table 7**.

### Differential Scanning Fluorimetry (DSF)

Purified P53 DBD constructs (Y220C and matching WT counterpart) encompassing amino acids 94-to −312 or 89-to-312 were diluted to a final concentration of 2 µM in a KH_2_PO_4_ buffer pH7.2 containing 150mM NaCl, 0.1mM TCEP and 0.0025% Tween20. Sypro dye (Invitrogen) was added to a 5X final concentration. DSF analysis was run on a Viia7 instrument (Thermo Fisher). The analysis method included an initial step of 3 minutes at 20°C, followed by a continuous ramp at 0.02°C/sec between 20°C and 65°C, and a final 1 minute step at 65°C. Experiments monitoring the effect of EDTA on the p53 DBD thermal stability included a pre-incubation of 10 minutes at 20°C with or without 5uM Rezatapopt followed by a 20 minute incubation at 20°C with or without 1mM EDTA. At the end of the incubation, DSF analysis was run using the method described above. For WT analysis with EDTA, overnight incubation was required for full stripping of the metal ion.

### FRET DNA binding assay

Oligonucleotides including p53 response elements were designed based on previous studies^21^. Sequences included palindromic 10 bp consensus p53 response elements (AGGCATGTCT) separated by 0 bp; 6 flanking base pairs were added 5’ to each response element half site. The detailed nucleotide sequences were as follow: forward p53 response element, 5’- CTG TCC AGG CAT GTC TAG ACA TGC CTA CAC CT - 3’; reverse p53 response element, 5’- AGG TGT AGG CAT GTC TAG ACA TGC CTG GAC AG - 3’. Oligonucleotide synthesis (Integrated DNA Technology, IDT) incorporated a biotin moiety at the 5’ end of the forward strand. All DNA strands were HPLC-purified. Oligonucleotide annealing used IDTE 1X TE solution, pH 8.0, and was performed according to the manufacturer recommendations. DNA binding ability of the Y220C mutant was probed following 28°C heat challenge. Briefly, the 5’-biotinylated duplex oligonucleotides were pre-bound for 20 minutes at 20°C to the FRET acceptor SA-d2 (1.5-fold excess relative to each DNA concentration) (Cisbio). p53 (94-312) or (89-312), WT or Y220C (35 nM), were pre-incubated with the DNA dose response (150 nM top concentration, 5-point dose response, 3-fold step dilutions) on ice for 20min in a buffer containing 50mM HEPES, pH 7.4, 150 mM NaCl, 0.1 mM TCEP, 0.01% Tween20, 0.1 mg/ml heat-denatured BSA (Biolabs). The protein/DNA complexes were then transferred to 20°C for 10 minutes before being submitted to 28°C heat challenge for 30 minutes. Following heat challenge, the protein/ DNA mixtures were returned on ice. The FRET donor Tb-FLAG-Antibody (0.2 nM) (Revvity) was added and incubated for 60 minutes before the FRET was read on a Perkin Elmer Envision instrument.

In experiments monitoring the Zn^2+^ binding affinity upon EDTA stripping at various temperatures, p53-DBD (89-312) (35nM) or P53 DBD-TD (89-360) (10nM), WT or Y220C, were mixed with DMSO or 5uM Rezatapopt, and incubated 1 hour with EDTA (4mM top concentration, 11-point dose response, 2-fold step dilutions) in a buffer containing 50mM HEPES, pH 7.4, 150 mM NaCl, 0.1mM TCEP, 0.01% Tween20, 0.1 mg/ml BSA at temperatures varying from 15°C to 32.5°C. At the end of each incubation, DNA oligonucleotides (300 nM) and FRET reagents (0.2 nM Tb-FLAG-Ab and 450nM SA-d2) were added and incubated 1 hour before the FRET was read on a Perkin Elmer Envision instrument. The dose-dependent rescue of Y220C Zn^2+^ affinity by Rezatapopt was evaluated following cross-titrations of the compound (5 uM top concentration, 6-point dose response, 2-fold step dilutions) and EDTA (4 mM top concentration, 11-point dose response, 2-fold step dilutions) at 20°C.

In experiments monitoring Rezatapopt potency in restoring Y220C DNA binding activity at 37°C, 10nM p53 DBD-TD (89-360) Y220C were pre-incubated at 20°C with compound dose-response (5 uM top concentration, 11-point dose response, 2-fold step dilutions) in a buffer containing 50 mM HEPES, pH 7.4, 150 mM NaCl, 0.1mM TCEP, 0.01% Tween20, 0.1 mg/ml BSA. Three hundred nanomolar oligonucleotides were then added and the protein/ compound/ DNA mixtures were incubated at 37°C for 10min to 90 min. At the end of each incubation, the mixtures were returned to 20°C for 10 minutes before the FRET reagents (0.2 nM Tb-FLAG-Ab and 450nM SA-d2) were added and incubated 1 hour before the FRET was read on a Perkin Elmer Envision instrument.

### FRET probe binding assay

The biotinylated probe binding affinity for the p53 DBD (89-312) Y220C was measured at temperatures varying from 4°C to 32.5°C. Briefly, 20nM p53 Y220C were incubated for 1 hour with **Compound 1** dose response (1.5 µM top concentration, 7-point dose response, 2-fold step dilutions) in a buffer containing 10mM KH_2_PO_4_, pH 7.2, 150mM NaCl, 0.1mM TCEP, 0.01% Tween20, 0.1 mg/ml BSA. FRET reagent donor (Tb-FLAG-Ab, 0.2 nM) and acceptor (SA-d2, 1.5-fold relative to each probe concentration) were pre-bound to the p53 Y220C and DNA oligonucleotides, respectively, when the temperatures of incubation remained below 30°C. For incubations running at temperatures exceeding 30°C, the FRET reagents were added to the plate after it was returned to 20°C and the FRET was read after 1 hour.

Rezatapopt binding affinity for the p53 DBD (89-312) Y220C at temperatures varying from 4°C to 32.5°C was evaluated from competition against the biotinylated probe FTX-0257987 set at EC_50_/K_d_ value for each temperature analyzed. Briefly, 20 nM p53 Y220C were incubated for 1 hour with **Compound 1** and a dose response of Rezatapopt (5 µM top concentration, 7-point dose response, 2-fold step dilutions) in a buffer containing 10 mM KH_2_PO_4_, pH 7.2, 150mM NaCl, 0.1 mM TCEP, 0.01% Tween20, 0.1 mg/ml BSA. As described above, the FRET reagents (Tb-FLAG-Ab, 0.2 nM; SA-d2, 1.5-fold relative to each probe concentration) were pre-bound for temperatures below 30°C or added to the plate after it was returned to 20°C for temperatures exceeding 30°C.

### Surface Plasmon Resonance Studies

SPR analysis was performed using an Cytiva S200 instrument at 20°C. C-terminal, Avi-tagged Y220C p53 (94-312) was immobilized to an SA chip up to ∼500RU using a running buffer consisting of 50mM potassium phosphate pH 7.2, 100mM NaCl, 0.01% Tween 20, 0.1mM TCEP, and 2% DMSO. Active surfaces were capped with pegylated biotin after immobilization. Reference surface was prepared by injecting pegylated biotin without any protein immobilized. Rezatapopt serial dilutions (top concentration of 500 nM; 2-fold dilution) were injected over the p53 surface at 30uL/min for 120sec with a 1500sec dissociation time. System was washed with 50% DMSO after each injection. Blank injections were included for double referencing and solvent correction was applied. Binding kinetic fits (1:1) were applied with refractive index parameter fixed to zero. Percent activity was calculated by dividing the fitted R_max_ for each dose response by the theoretical R_max_ derived from ligand and analyte molecular weights.

### Unbiased Molecular Dynamics Simulations

Molecular dynamics (MD) simulations were performed using Cresset Flare v10 software, utilizing the Open Force Field (OpenFF) 2.2.0 with explicit TIP4Pew water molecules. The total duration of each simulation was 200 ns. The systems were solvated in a truncated octahedral solvent box with a solvent buffer of 10 Å. Solvent ionic strength was adjusted by adding counterions to achieve a neutral charge. Simulations were run under isothermal-isobaric (NPT) conditions at a temperature of 298 K and pressure of 1.0 bar. Non-bonded interactions were treated automatically with a cutoff of 12 Å. Energy minimization and equilibration were performed automatically for 200 ps using standard protocols prior to production runs. The integration timestep for all MD simulations was set to 4 fs, applying hydrogen mass repartitioning with a scaling factor of 1.50. Trajectories were recorded by saving every 20 ps, resulting in 10,000 frames over the 200 ns simulation interval.

### Adaptively Biased Molecular Dynamics Simulations

Biased simulations were first equilibrated using GROMACS 5.0.5. All simulations were solvated in TIP3P water and salted (NaCl) to 150 mM. The nearest box face was at least 1nm from the protein. All simulations used the CHARMM27 force field within GROMACS. Initial atomic coordinates were first energy minimized to a maximum force of 1000 kJ/mol/nm. Next, 5000 steps in the canonical ensemble at 300 Kelvin, with heavy atoms restrained, were used to further relax the system. Finally, 5 nanoseconds in the isothermal isobaric ensemble were used to bring the system to atmospheric pressure. All production simulations were 2 microseconds long and used the adaptive biasing scheme of fABMACS^15^. Overfill protection^16^ was used to limit the flood level to 10 kJ/mol for each biased atom. In these simulations each biased atom had its own 3-dimensional depth-limited bias potential. 15 atoms, identical across simulations, were biased. The position and total bias potential energy were written every 50000 steps and used in Diffusion Maps^17^ for dimensionality reduction. The diffusion map projections were rewieghted^22^ to generate a free energy estimate on the diffusion map. Biased atoms are shown in **Table 2**

**Table 2:**
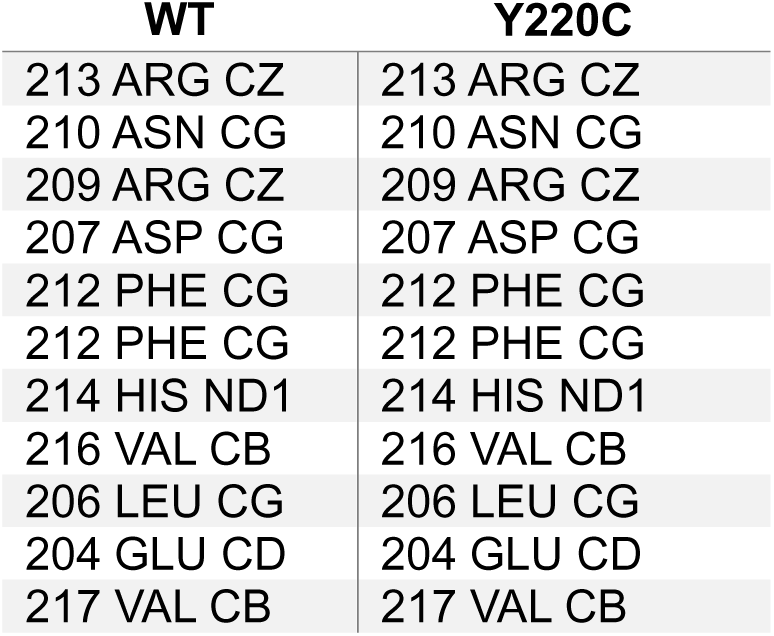
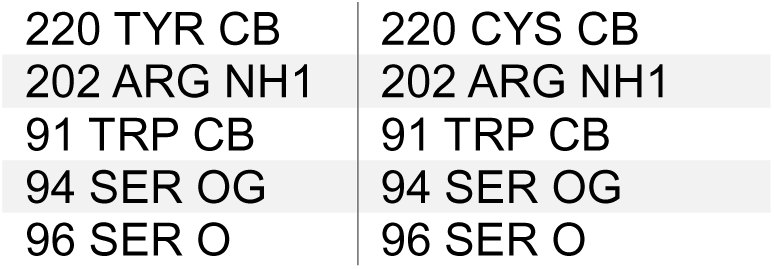
Biased Atoms for Adaptively Biased Molecular Dynamics Simulations.

### Hydrogen-Deuterium Exchanges Studies (HDX)

For stabilization of Y220C by Rezatapopt, 20 µM Y220C (89-312-GGSGGS-FLAG) was treated with 40 µM Rezatapopt or DMSO (0.4%) for 15 minutes at 20°C in 50 mM K_2_HPO_4_, pH 7.2, 100 mM NaCl. Mixtures were then incubated at 30°C for 1, 30 or 60 minutes minutes. After the respective incubations, samples were cooled at 20°C for 2 minutes. One micoliter of protein mixture was transferred to 18 µL of labeling buffer 50 mM K_2_HPO_4_, pH 7.2, 100 mM NaCl in D_2_O at 20°C. Labeling was performed for 10 seconds. Labeling was quenched trhough addition of 19 mL (1:1) of 150 mM KH_2_PO_4_, pH 2.485 (f.c. 0.5 µM Y220C). Samples were flash frozen until analysis. All 38 µL of samples were injected for analysis.

**Table 3:**
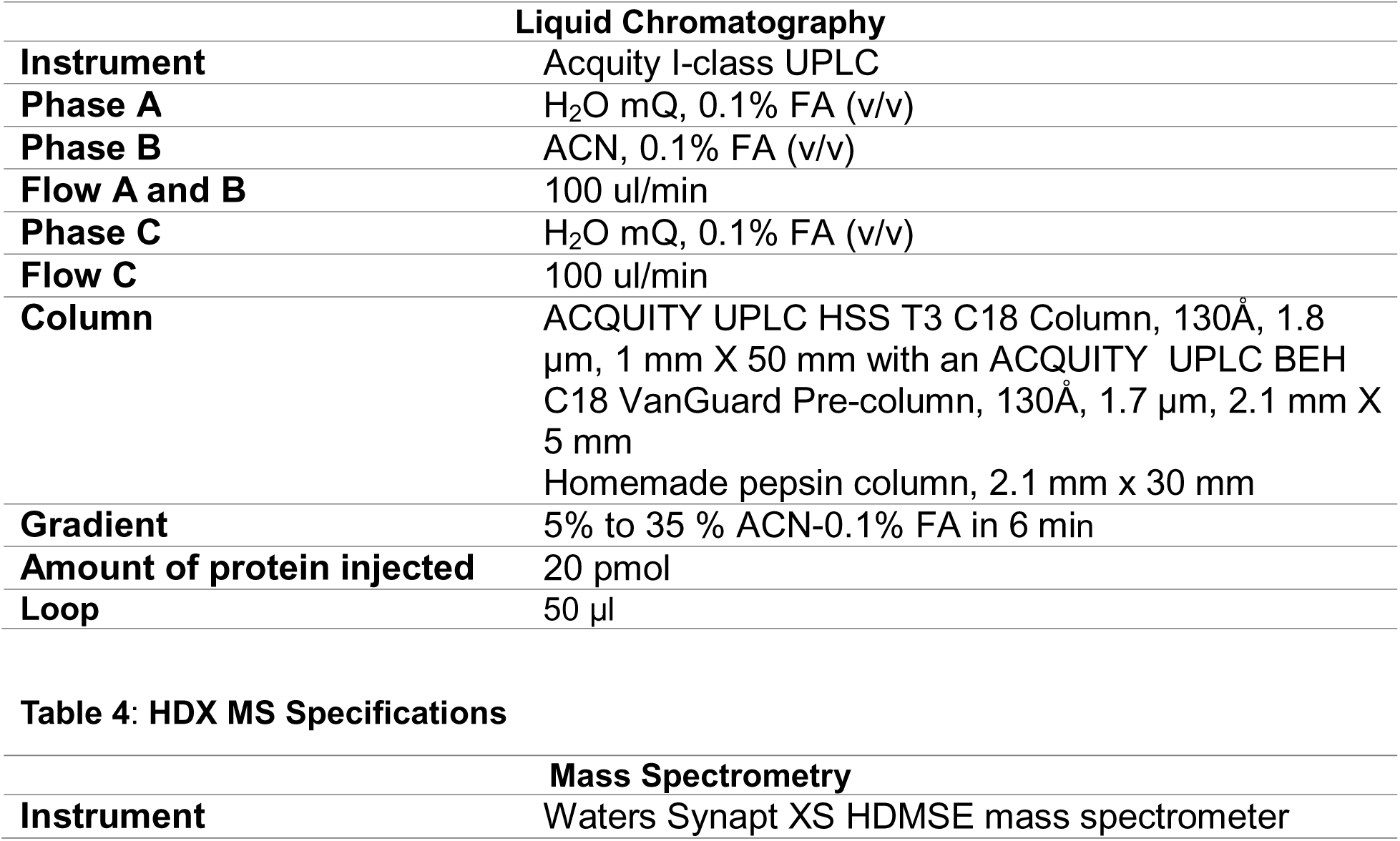
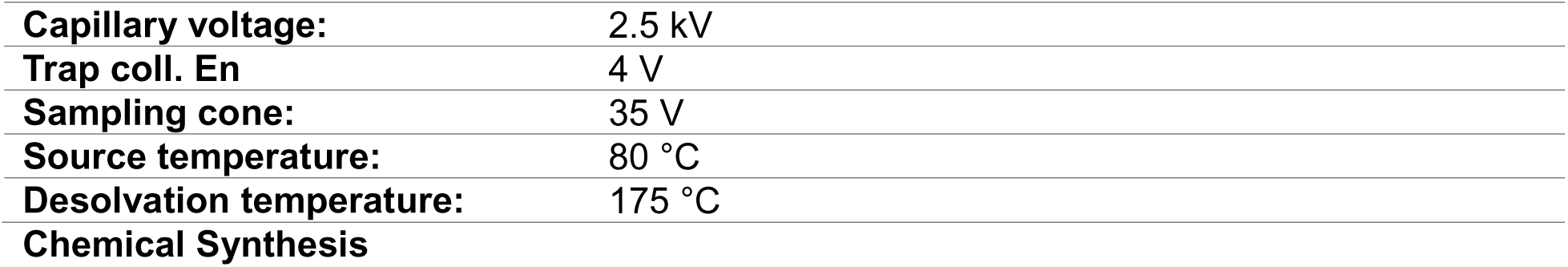
HDX LC Specifications.

Rezatapopt, Compound 2 and Compound 3 were synthesized in accordance with previous disclosures^4,23–26^

Preparation of Y220C TP53 TR-FRET binding probe is described in the Supporting Information

## References

1. Joerger, A. C., Ang, H. C. & Fersht, A. R. Structural basis for understanding oncogenic p53 mutations and designing rescue drugs. Proc National Acad Sci 103, 15056–15061 (2006).

2. Liu, X. et al. Small molecule induced reactivation of mutant p53 in cancer cells. Nucleic Acids Res 41, 6034–6044 (2013).

3. Bauer, M. R. et al. A structure-guided molecular chaperone approach for restoring the transcriptional activity of the p53 cancer mutant Y220C. Future Med Chem 11, 2491–2504 (2019).

4. Vu, B. T. et al. Discovery of Rezatapopt (PC14586), a First-in-Class, Small-Molecule Reactivator of p53 Y220C Mutant in Development. ACS Med. Chem. Lett. 16, 34–39 (2025).

5. Puzio-Kuter, A. M. et al. Restoration of the Tumor Suppressor Function of Y220C-Mutant p53 by Rezatapopt, a Small-Molecule Reactivator. Cancer Discov. 15, 1159–1179 (2025).

6. Boeckler, F. M. et al. Targeted rescue of a destabilized mutant of p53 by an in silico screened drug. Proc National Acad Sci 105, 10360–10365 (2008).

7. Guiley, K. Z. & Shokat, K. M. A small molecule reacts with the p53 somatic mutant Y220C to rescue wild-type thermal stability. Cancer Discov (2022) doi:10.1158/2159-8290.cd-22-0381.

8. Baud, M. G. J. et al. Aminobenzothiazole derivatives stabilize the thermolabile p53 cancer mutant Y220C and show anticancer activity in p53-Y220C cell lines. Eur J Med Chem 152, 101–114 (2018).

9. Bullock, A. N., Henckel, J. & Fersht, A. R. Quantitative analysis of residual folding and DNA binding in mutant p53 core domain: definition of mutant states for rescue in cancer therapy. Oncogene 19, 1245–1256 (2000).

10. Blanden, A. R. et al. Zinc shapes the folding landscape of p53 and establishes a pathway for reactivating structurally diverse cancer mutants. Elife 9, e61487 (2020).

11. Joerger, A. C., Ang, H. C., Veprintsev, D. B., Blair, C. M. & Fersht, A. R. Structures of p53 Cancer Mutants and Mechanism of Rescue by Second-site Suppressor Mutations*. J. Biol. Chem. 280, 16030–16037 (2005).

12. Natan, E. et al. Interaction of the p53 DNA-Binding Domain with Its N-Terminal Extension Modulates the Stability of the p53 Tetramer. J Mol Biol 409, 358–368 (2011).

13. Bauer, M. R. et al. Targeting Cavity-Creating p53 Cancer Mutations with Small-Molecule Stabilizers: the Y220X Paradigm. Acs Chem Biol 15, 657–668 (2020).

14. Joerger, A. C., Ang, H. C., Veprintsev, D. B., Blair, C. M. & Fersht, A. R. Structures of p53 Cancer Mutants and Mechanism of Rescue by Second-site Suppressor Mutations*. J. Biol. Chem. 280, 16030–16037 (2005).

15. Dickson, B. M., Waal, P. W. de, Ramjan, Z. H., Xu, H. E. & Rothbart, S. B. A fast, open source implementation of adaptive biasing potentials uncovers a ligand design strategy for the chromatin regulator BRD4. J. Chem. Phys. 145, 154113 (2016).

16. Dickson, B. M. Overfill Protection and Hyperdynamics in Adaptively Biased Simulations. J. Chem. Theory Comput. 13, 5925–5932 (2017).

17. Coifman, R. R. et al. Geometric diffusions as a tool for harmonic analysis and structure definition of data: Diffusion maps. Proc. Natl. Acad. Sci. 102, 7426–7431 (2005).

18. Yung-Chi, C. & Prusoff, W. H. Relationship between the inhibition constant (KI) and the concentration of inhibitor which causes 50 per cent inhibition (I50) of an enzymatic reaction. Biochem. Pharmacol. 22, 3099–3108 (1973).

19. Kabsch, W. XDS. Acta Crystallogr. Sect. D 66, 125–132 (2010).

20. Agirre, J. et al. The CCP4 suite: integrative software for macromolecular crystallography. Acta Crystallogr. Sect. D 79, 449–461 (2023).

21. El-Deiry, W. S., Kern, S. E., Pietenpol, J. A., Kinzler, K. W. & Vogelstein, B. Definition of a consensus binding site for p53. Nat Genet 1, 45–49 (1992).

22. Schäfer, T. M. & Settanni, G. Data Reweighting in Metadynamics Simulations. J. Chem. Theory Comput. 16, 2042–2052 (2020).

23. Binh, V., Romyr, D., Hongju, L., Bruce, F. & Andrew, G. METHODS AND COMPOUNDS FOR RESTORING MUTANT p53 FUNCTION. (2021).

24. Vu et al. METHODS AND COMPOUNDS FOR RESTORING MUTANT p53 FUNCTION.

25. Vu et al. METHODS AND COMPOUNDS FOR RESTORING MUTANT p53 FUNCTION.

26. Arnold, L., David, M., Binh, V., W, D. T. & Melissa, D. METHODS AND COMPOUNDS FOR RESTORING MUTANT p53 FUNCTION. (2021).

