## Supplemental Information for "Thermodynamic Insights into the WT and Y220C TP53 DBDs Reveals that the Oncogenic Y220C Variant is a Loss of Function Mutation for Zn^2+^-binding at Physiological Temperature"

| Supplementary Figure 1 | *Design and Characterization of an Endogenous TP53 Y220C DBD Construct.* | p. S2 |
| --- | --- | --- |
| Supplementary Figure 2 | *Temperature-Dependent Destabilization of Zn^2+^ Binding to the p53 DBD.* | p. S2 |
| Supplementary Figure 3. | *Temperature-Dependent restoration of Zn^2+^ binding induced by Rezatapopt to the Y220C p53 DBD.* | p. S3 |
| Supplementary Figure 4. | *Development and Validation of Binding Affinity Assays for Thermodynamic Characterization of Rezatapopt Binding.* | p. S3 |
| Supplementary Figure 5. | *Temperature-Dependent Binding Studies through probe Displacement of Rezatapopt and Related Analogs* | p. S4 |
| Supplementary Figure 6. | *Overlay of Structure of Compound 3 and Resulting Clash with Representative Frame MD Frame of* ${Y220C}_{apo}^{II}$*.* | p. S4 |
| Supplementary Table 1. | *EDTA IC_50_ for WT and Y220C Constructs at Various Temperatures.* | p. S5 |
| Supplementary Table 2. | *Stabilization of Zn-Binding by Rezatapopt to Y220C* | p. S5 |
| Supplementary Table 3. | *Restoration of WT-level Zn^2+^ affinity to Y220C by Rezatapopt at Various Temperatures.* | p. S5 |
| Supplementary Table 4. | *Rezatapopt EC_50_ over Time at 37ºC for Restoration of DNA Binding to the Y220C DBD-TD* | p. S6 |
| Supplementary Table 5. | *Summary of Temperature-Dependent Binding Studies* | p. S7 |
| Supplementary Table 6. | *Crystallographic Data Collection and Refinement Statistics* | p. S8 |
| Supplementary Table 7. | *Crystallographic Data Collection and Refinement Statistics* | p. S9 |
| Chemical Synthesis | *Preparation of Y220C TP53 TR-FRET binding probe* | pp. S10-S15 |

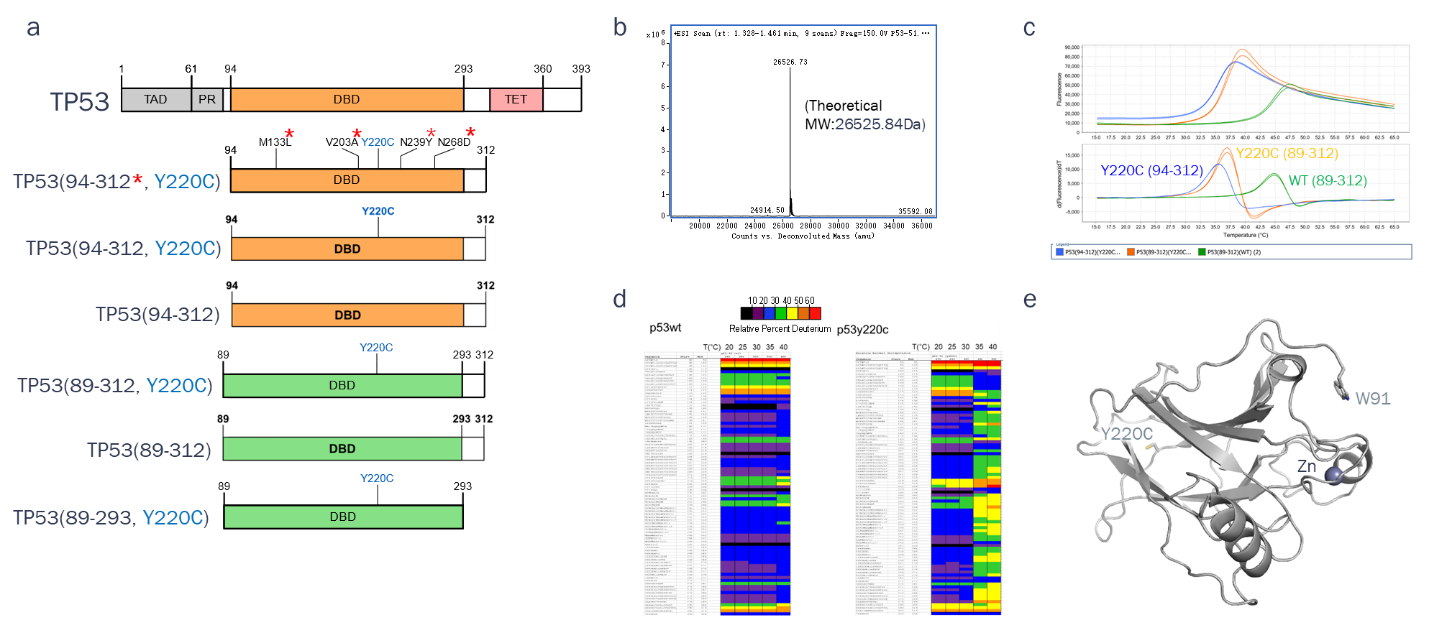

**Supplementary** **Figure 1. Design and Characterization of an Endogenous TP53 Y220C DBD Construct.** (**a**) Domain architecture of TP53 and Y220C construct. (**b**) Intact MS spectrum for endogenous Y220C DBD (res. 89-312). (**c**) DSF raw fluorescence (top) and derivative traces (bottom) of TP53 Y220C (94-312), TP53 Y220C (89-312) and TP53 WT (89-312). Duplicate measurements were taken. T_m_ values were determined to be 36.56 ºC ± 0ºC for TP53 Y220C (94-312), 36.96 ºC ± 0ºC for TP53 Y220C (89-312) and 44.83 ºC ± 0ºC TP53 WT (89-312). (**d**) Deuterium uptake plots for TP53 Y220C (89-312) and TP53 WT (89-312). Data represent the mean of duplicate experiments where protein constructs were subjected to the reported temperatures for 10 minutes before cooling back to room temperature (RT). Labeling with D_2_O was then performed for 10 seconds. (**e**) Cartoon depiction of TP53 Y220C (89-293).

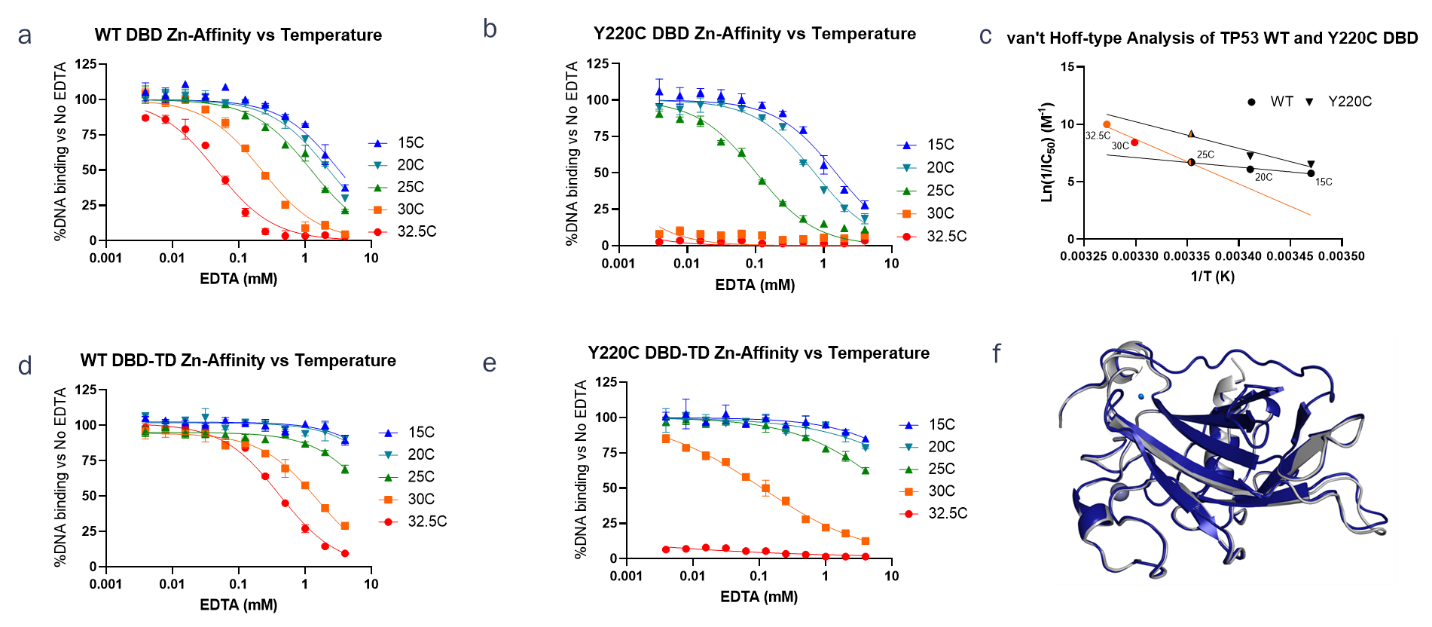

**Supplementary Figure 2. Temperature-Dependent Destabilization of Zn^2+^ Binding to the p53 DBD.** IC_50_ Values are summarized in **Supplementary Table 1.**

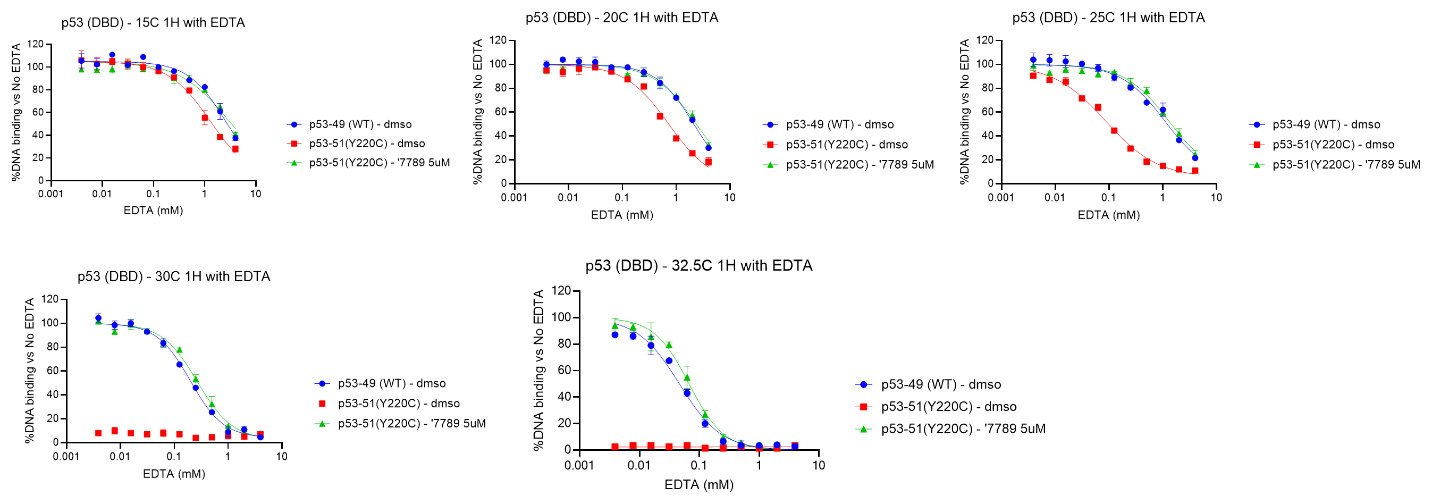

**Supplementary Figure 3**. **Temperature-Dependent restoration of Zn^2+^ binding induced by Rezatapopt to the Y220C p53 DBD.** IC_50_ Values are summarized in **Supplementary Table 3.**

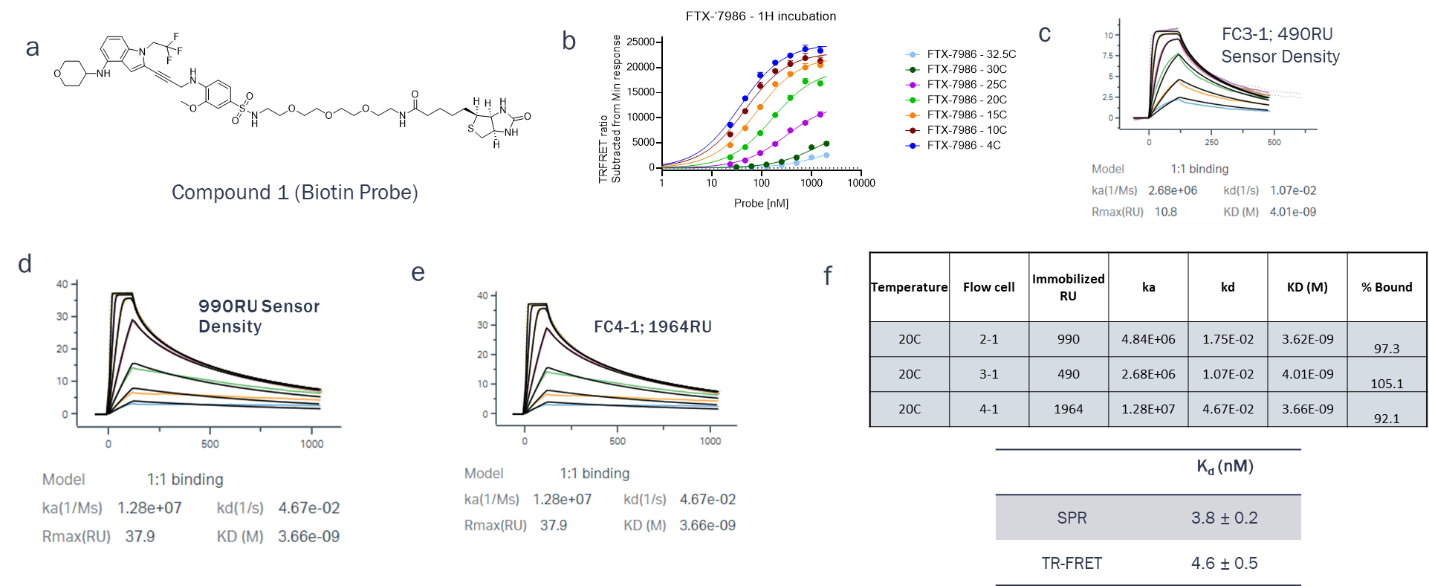

**Supplementary Figure 4. Development and Validation of Binding Affinity Assays for Thermodynamic Characterization of Rezatapopt Binding.**

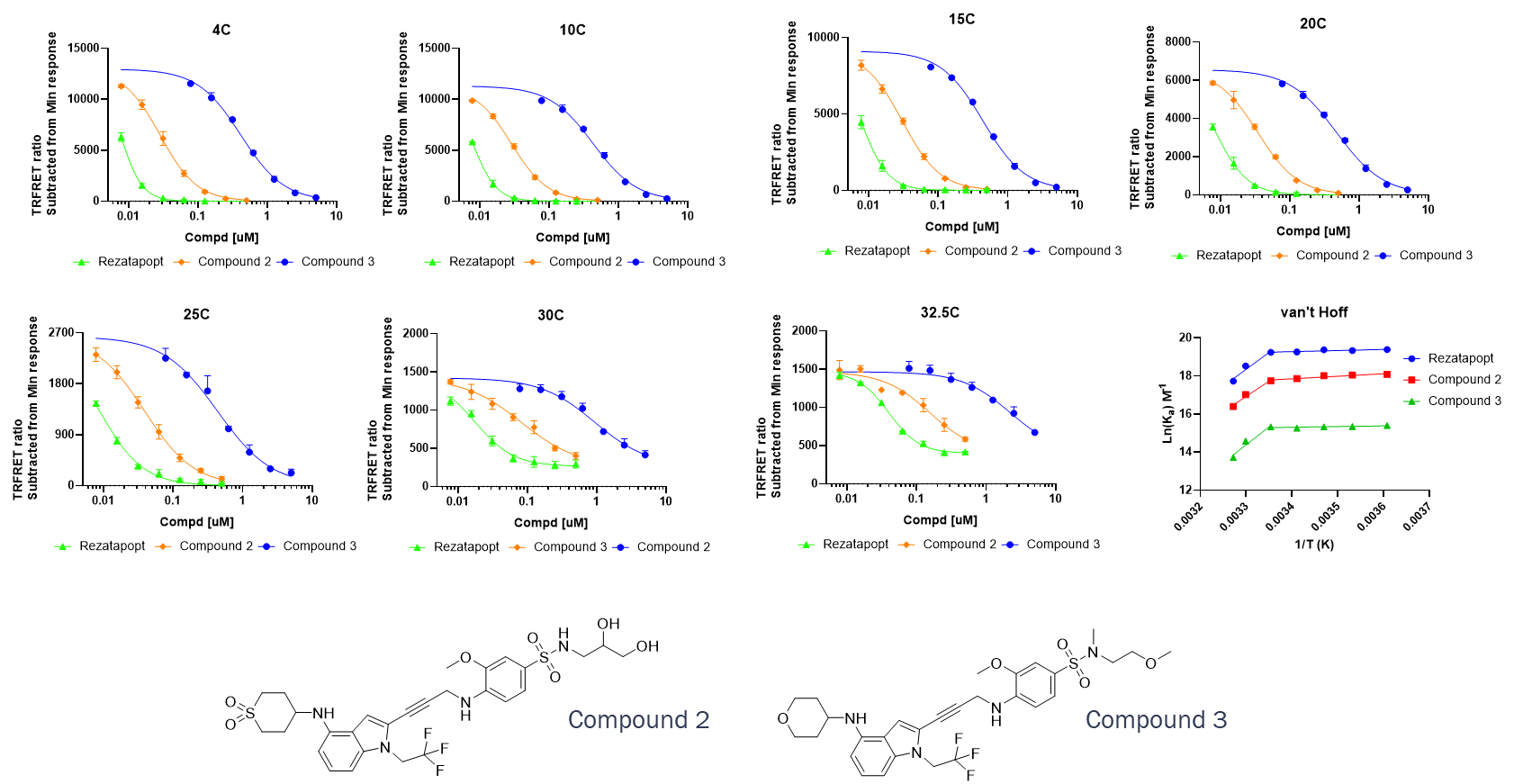

**Supplementary Figure 5. Temperature-Dependent Binding Studies through Probe Displacement of Rezatapopt and Related Analogs**

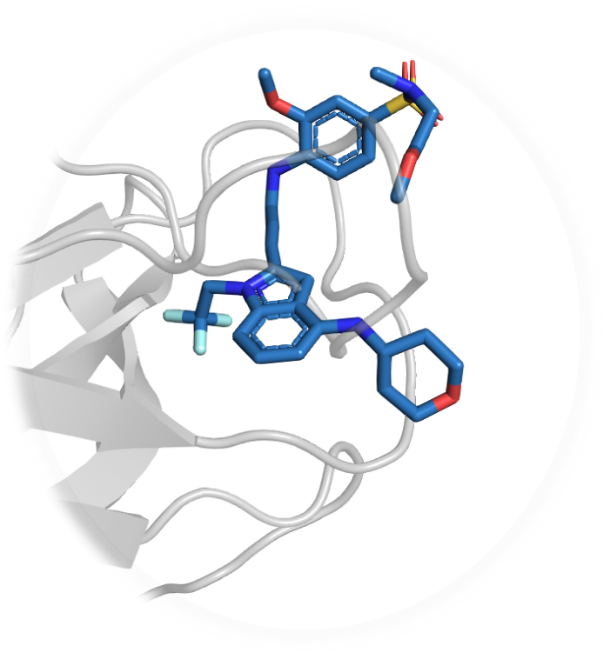

**Supplementary Figure 6. Overlay of Structure of Compound 3 and Resulting Clash with Representative Frame MD Frame of** ${\boldsymbol{Y}\boldsymbol{220}\boldsymbol{C}}_{\boldsymbol{apo}}^{\boldsymbol{II}}$**.**

**Supplementary Table 1. EDTA IC_50_ for WT and Y220C Constructs at Various Temperatures.**

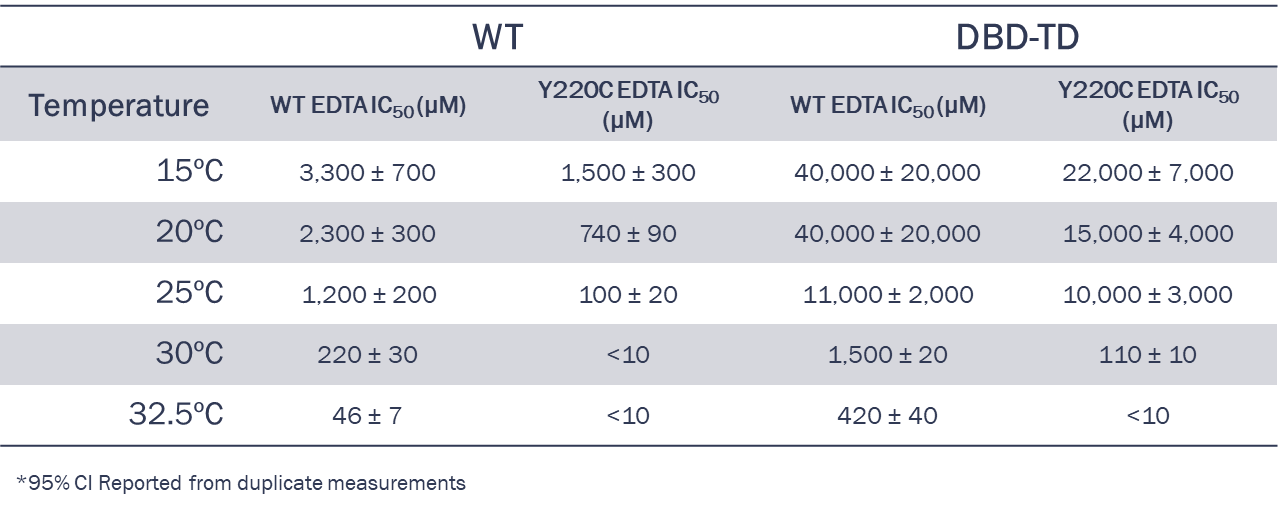

**Supplementary Table 2. Stabilization of Zn-Binding by Rezatapopt to Y220C**

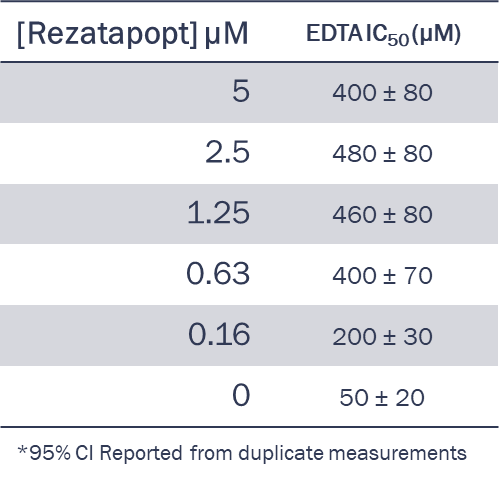

**Supplementary Table 3. Restoration of WT-level Zn^2+^ affinity to Y220C by Rezatapopt at Various Temperatures.**

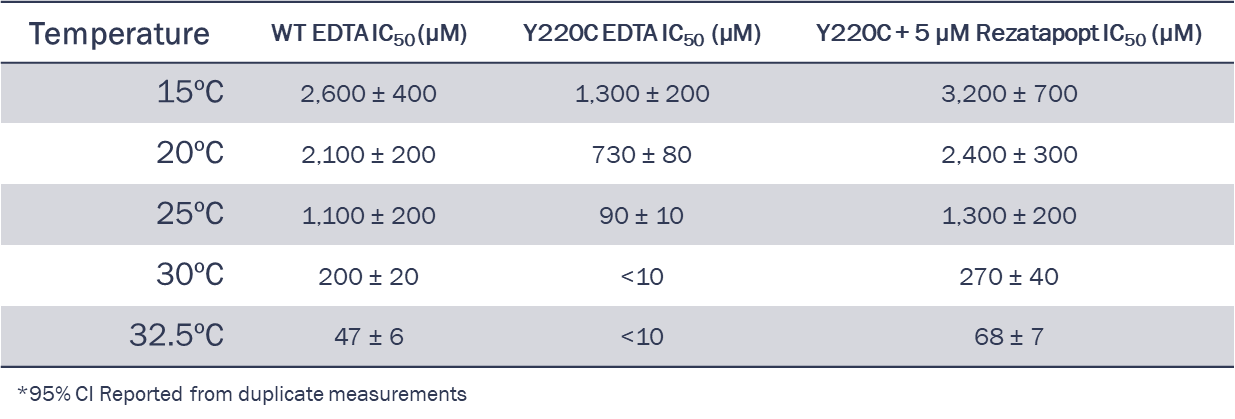

**Supplementary Table 4. Rezatapopt EC_50_ over Time at 37ºC for Restoration of DNA Binding to the Y220C DBD-TD**

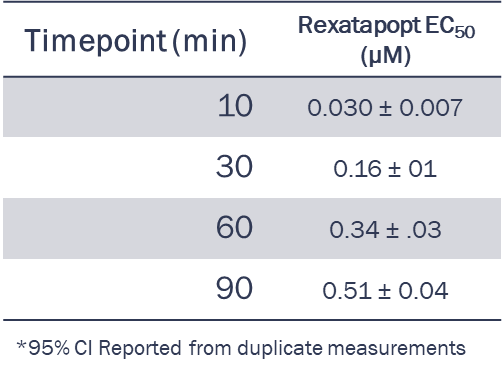

**Supplementary Table 5. Summary of Temperature-Dependent Binding Studies**

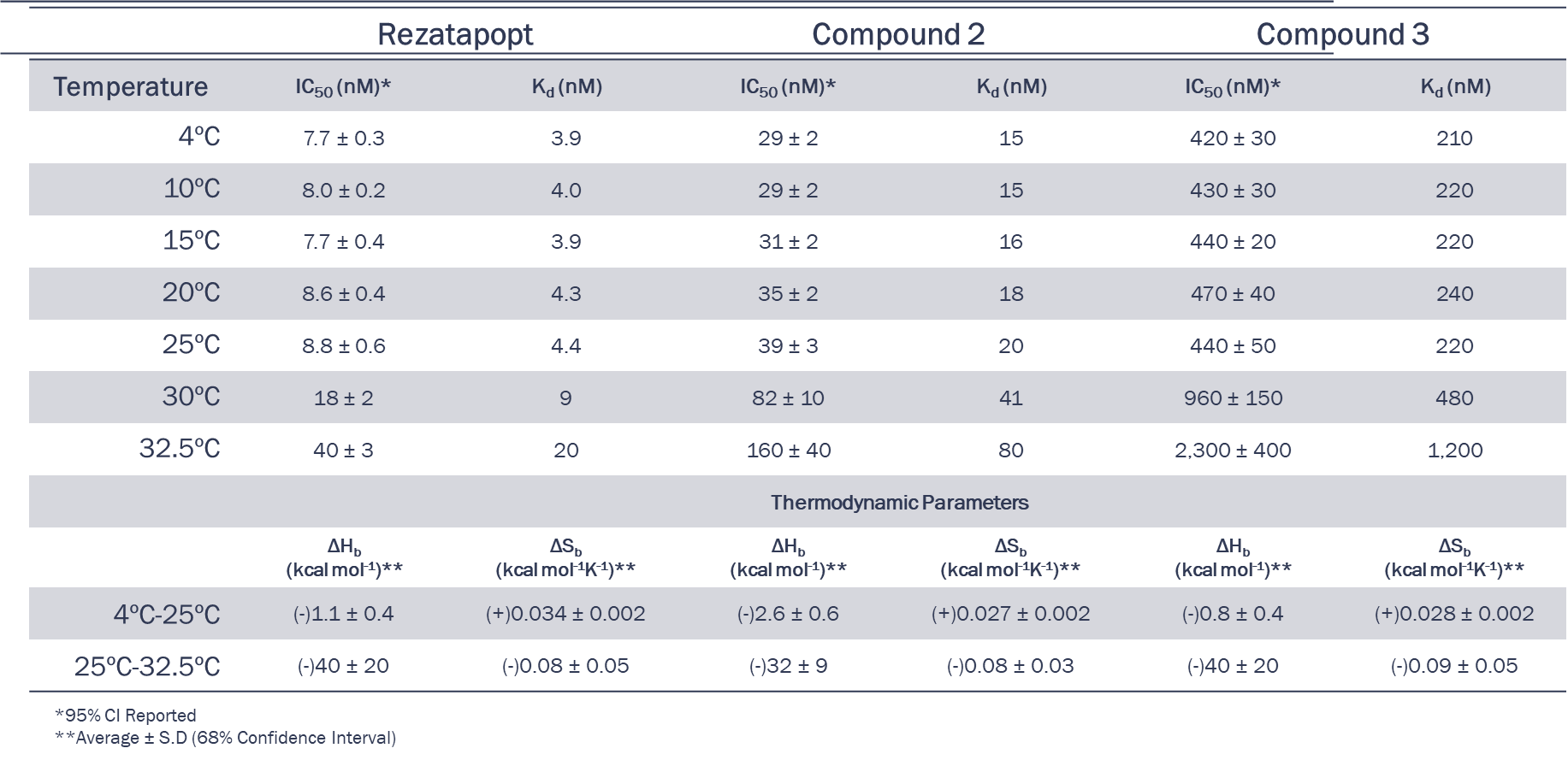

**Supplementary Table 6. Crystallographic Data Collection and Refinement Statistics**

|  | TP53 (89-293, Y220C) |
| --- | --- |
| **Data Collection** |  |
| Space group | *P2_1_* |
| Cell dimensions |  |
| *a, b, c* (Å) | 50.31 68.44 60.12 |
| α, β, γ (°) | 90.00 98.81 90.00 |
| Resolution (Å) | 44.87 – 2.11 (2.17 – 2.11) |
| R*merge* | 0.113 (0.830) |
| R*meas* | 0.134 (0.985) |
| R*pim* | 0.071 (0.526) |
| *I* / σ*I* | 11.0 (2.3) |
| *CC*1/2 | 0.997 (0.811) |
| Completeness (%) | 99.9 (100.0) |
| Redundancy | 6.7 (6.7) |
| **Refinement** |  |
| Resolution (Å) | 44.87 – 2.11 (2.17 – 2.11) |
| No. reflections | 22142 |
| *Rwork* / *Rfree* | 0.19259/0.26688 |
| No. atoms | 3299 |
| Protein | 3167 |
| Ligand | 0 |
| Water  Zn | 130  2 |
| *B*-factors | 40.25 |
| Protein | 43.56 |
| Ligand | N/A |
| Water  Zn | 39.92  38.35 |
| R.m.s. deviations |  |
| Bond lengths (Å) | 0.007 |
| Bond angles (°) | 1.493 |
| Ramachandran plot |  |
| Favored (%) | 95.1 |
| Allowed (%) | 4.9 |
| Outliers (%) | 0 |
| Statistics for the highest-resolution shell are shown in parentheses. |  |

**Supplementary Table 7. Crystallographic Data Collection and Refinement Statistics**

|  | TP53 (89-293, Y220C) +  Compound 3 |
| --- | --- |
| **Data Collection** |  |
| Space group | *P2_1_* |
| Cell dimensions |  |
| *a, b, c* (Å) | 50.49 68.54 59.60 |
| α, β, γ (°) | 90.00 99.27 90.00 |
| Resolution (Å) | 44.64 – 2.39 (2.48 – 2.39) |
| R*merge* | 0.103 (0.892) |
| R*meas* | 0.122 (1.062) |
| R*pim* | 0.065 (0.570) |
| *I* / σ*I* | 10.7 (2.2) |
| *CC*1/2 | 0.996 (0.814) |
| Completeness (%) | 99.9 (100.0) |
| Redundancy | 6.8 (6.7) |
| **Refinement** |  |
| Resolution (Å) | 44.64 – 2.39 (2.45 – 2.39) |
| No. reflections | 15187 |
| *Rwork* / *Rfree* | 0.21373/0.29768 |
| No. atoms | 3241 |
| Protein | 3131 |
| Ligand | 84 |
| Water  Zn | 24  2 |
| *B*-factors | 58.47 |
| Protein | 87.08 |
| Ligand | 67.48 |
| Water  Zn | 54.01  70.66 |
| R.m.s. deviations |  |
| Bond lengths (Å) | 0.006 |
| Bond angles (°) | 1.403 |
| Ramachandran plot |  |
| Favored (%) | 90.0 |
| Allowed (%) | 8.0 |
| Outliers (%) | 2.0 |
| Statistics for the highest-resolution shell are shown in parentheses. |  |

**Chemical Synthesis**

*Preparation of Y220C TP53 TR-FRET binding probe*

BP1, 5-[(3aS,4S,6aR)-2-oxo-1,3,3a,4,6,6a-hexahydrothieno[3,4-d]imidazol-4-yl]-N-[2-[2-[2-[2-[[3-methoxy-4-[3-[4-(tetrahydropyran-4-ylamino)-1-(2,2,2-trifluoroethyl)indol-2-yl]prop-2-ynylamino]phenyl]sulfonylamino]ethoxy]ethoxy]ethoxy]ethyl]pentanamide

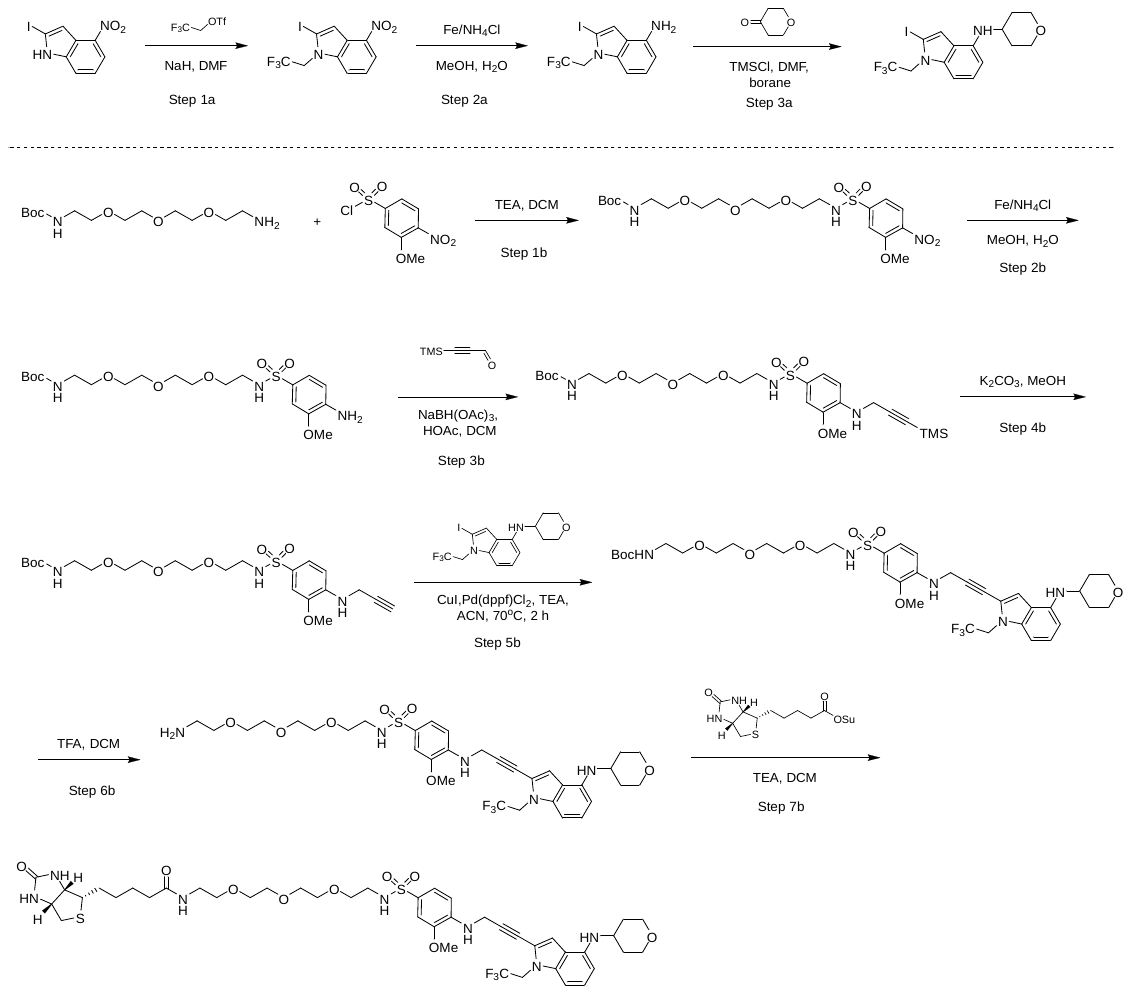

BP1 synthesis, step 1a, 2-iodo-4-nitro-1-(2,2,2-trifluoroethyl)indole: To a solution of 2-iodo-4-nitro-1H-indole (3.0 g, 10.4 mmol, 1.0 equiv.) in THF (20 mL) was added NaH (2.0 g, 52.0 mmol, 5.0 equiv.; 60.0% dispersion in oil) at 0°C by portions, and stirred at 0°C for 30 minutes. 2,2,2-trifluoroethyl trifluoromethanesulfonate (9.6 g, 41.0 mmol, 4.0 equiv.) was added to the reaction mixture at 0°C in portions. The mixture was stirred at 25°C for 2 hours. The reaction mixture was quenched with H_2_O (20 mL) and the resulting mixture was partitioned between EtOAc (200 mL) and H_2_O (200 mL) and the aqueous layer was extracted with EtOAc (2 × 200 mL). The combined organic layers were washed with brine (50 mL), dried over anhydrous Na_2_SO_4_, filtered, and concentrated under reduced pressure. The resulting residue was purified by silica gel column chromatography (15:1 to 12:1 petroleum ether:EtOAc) to afford the title compound (5.0 g, crude) as yellow solid.

1H NMR (400 MHz, CHLOROFORM-d) δ 8.15 (d, J = 8.2 Hz, 1H), 7.80 - 7.63 (m, 2H), 7.33 (t, J = 8.2 Hz, 1H), 4.85 (q, J = 8.2 Hz, 2H).

BP1 synthesis, step 2a, 2-iodo-1-(2,2,2-trifluoroethyl)indol-4-amine: To a solution of 2-iodo-4-nitro-1-(2,2,2-trifluoroethyl)indole (1.9 g, 5.1 mmol, 1.0 equiv.) in EtOH (20.0 mL) and H_2_O (5.0 mL) was added Fe (717 mg, 12.8 mmol, 2.5 equiv.) and NH_4_Cl (687 mg, 12.8 mmol, 2.5 equiv.). The mixture was stirred at 80 °C for 2 hours. The reaction solution was filtered through a pad of diatomite and the filtrate was partitioned between EtOAc (200 mL) and H_2_O (200 mL). The aqueous layer was extracted with EtOAc (2 × 200 mL). The combined organic layers were washed with brine (50 mL), dried over anhydrous Na_2_SO_4_, filtered, and concentrated under reduced pressure to afford the title compound (1.7 g, 97% yield) as yellow solid.

LCMS [M+1] = 341.1.

BP1 synthesis, step 3a, 2-iodo-N-tetrahydropyran-4-yl-1-(2,2,2-trifluoroethyl)indol-4-amine: To a solution of 2-iodo-1-(2,2,2-trifluoroethyl)indol-4-amine (600 mg, 1.7 mmol, 1.0 equiv.) in DMF (10 mL) was added chloro(trimethyl)silane (559.8 μL, 4.4 mmol, 2.5 equiv.) and tetrahydropyran-4-one (648.1 μL, 7.0 mmol, 4.0 equiv.). The mixture was stirred at 0°C for 2 hours. Borane-tetrahydrofuran complex (1 M, 8.8 mL, 5.0 equiv.) was added to the mixture under N_2_ and the resulting mixture was stirred at 0-20°C for 12 hours. The reaction was poured into a saturated aqueous solution of NH_4_Cl (1.5 ml) and extracted with EtOAc (3 × 5 mL), dried over Na_2_SO_4_, filtered and concentrated under reduced pressure to afford the title compound (600 mg, 80% yield) as white solid.

^1^H NMR (400 MHz, CHLOROFORM-*d*) δ 8.03 (s, 1H), 7.07 (t, *J* = 8.0 Hz, 1H), 6.90 - 6.70 (m, 2H), 6.34 (d, *J* = 7.8 Hz, 1H), 4.69 (q, *J* = 8.4 Hz, 2H), 4.11 - 4.00 (m, 2H), 3.92 - 3.71 (m, 1H), 3.73 - 3.63 (m, 1H), 3.56 (t, *J* = 10.6 Hz, 2H), 2.97 (s, 2H), 2.89 (s, 1H), 2.68 (d, *J* = 9.4 Hz, 1H), 2.24 - 2.01 (m, 2H).

LCMS [M+1] = 425.1.

BP1 synthesis, step 1b, *tert*-Butyl N-[2-[2-[2-[2-[(3-methoxy-4-nitro-phenyl)sulfonylamino]ethoxy]ethoxy]-ethoxy]ethyl]carbamate: To a solution of *tert*-butyl N-[2-[2-[2-(2-aminoethoxy)ethoxy]ethoxy]-ethyl]carbamate (906 mg, 3.1 mmol, 1.3 equiv.) in DCM (15 mL) was added TEA (1.6 mL, 11.9 mmol, 5.0 equiv.) and 3-methoxy-4-nitro-benzenesulfonyl chloride (600 mg, 2.4 mmol, 1.0 equiv.). The mixture was stirred at 15°C for 1 hour. The reaction mixture was concentrated under reduced pressure and the resulting residue was partitioned between EtOAc (50 mL) and H_2_O (30 mL) and the aqueous layer was extracted with EtOAc (2 × 50 mL). The combined organic layers were washed with brine (50 mL), dried over anhydrous Na_2_SO_4_, filtered, and concentrated under reduced pressure to afford the title compound (1.1 g, 91% yield) as a brown oil.

^1^H NMR (400 MHz, DMSO-*d*_6_) δ 8.07 (d, *J* = 8.4 Hz, 1H), 7.67 (s, 1H), 7.51 (d, J = 8.4 Hz, 1H), 6.75 (s, 1H), 4.00 (s, 3H), 3.53 - 3.41 (m, 10H), 3.15 - 2.90 (m, 6H), 2.65 (t, *J* = 5.7 Hz, 1H), 1.36 (s, 9H)

LCMS [M+1] = 408.2.

BP1 synthesis, step 2b, *tert*-Butyl N-[2-[2-[2-[2-[(4-amino-3-methoxy-phenyl)sulfonylamino]ethoxy]ethoxy]-ethoxy]ethyl]carbamate: To a solution of *tert*-butyl N-[2-[2-[2-[2-[(3-methoxy-4-nitro-phenyl)sulfonylamino]ethoxy]ethoxy]ethoxy]-ethyl]carbamate (2.0 g, 3.9 mmol, 1.0 equiv.) in EtOH (12.0 mL) and H_2_O (3.0 mL) was added Fe (1.1 g, 19.7 mmol, 5.0 equiv.) and NH_4_Cl (1.1 g, 19.7 mmol, 5.0 equiv.). The mixture was stirred at 80°C for 1 hour. The suspension was filtered through a pad of Celite and the pad cake was washed with EtOH (3 × 20 mL). The filtrate was concentrated under reduced pressure and the resulting residue was partitioned between EtOAc (50 mL) and H_2_O (30 mL). Then the aqueous layer was extracted again with EtOAc (2 × 50 mL). The combined organic layers were washed with brine (50 mL), dried over anhydrous Na_2_SO_4_, filtered, and concentrated under reduced pressure to afford the title compound (1.5 g, 80% yield) as a brown oil.

^1^H NMR (400 MHz, DMSO-*d*_6_) δ 7.20 (t, *J* = 6.0 Hz, 1H), 7.16 - 7.13 (m, 1H), 7.12 (s, 1H), 6.76 (br t, *J* = 5.6 Hz, 1H), 6.69 - 6.63 (m, 1H), 6.73 - 6.60 (m, 1H), 5.56 (s, 2H), 3.80 (s, 3H), 3.50 - 3.42 (m, 7H), 3.40 - 3.35 (m, 3H), 3.05 (q, *J* = 6.0 Hz, 2H), 2.80 (q, *J* = 6.0 Hz, 2H), 1.36 (s, 9H)

LCMS [M+1] = 378.3.

BP1 synthesis, step 3b, *tert*-Butyl N-[2-[2-[2-[2-[[3-methoxy-4-(3-trimethylsilylprop-2-ynylamino)phenyl]-sulfonylamino]ethoxy]ethoxy]ethoxy]ethyl]carbamate: To a solution of *tert*-butyl N-[2-[2-[2-[2-[(4-amino-3-methoxy-phenyl)sulfonylamino]ethoxy]ethoxy]ethoxy]-ethyl]carbamate (1.5 g, 3.1 mmol, 1.0 equiv.) in DCM (15 mL) and acetic acid (3 mL) was added 3-trimethylsilylprop-2-ynal (396 mg, 3.1 mmol, 1.0 equiv.). The mixture was stirred at 35°C for 17 hours and then sodium triacetoxyborohydride (2.6 g, 12.5 mmol, 4.0 equiv.) was added to the mixture. The mixture was stirred at 35°C for 17 hours. The reaction mixture was diluted with EtOAc (50 mL) and H_2_O (30 mL) and the aqueous layer was extracted with EtOAc (2 × 50 ml). The combined organic extracts were washed with brine (50 mL), dried over anhydrous Na_2_SO_4_, filtered, and concentrated under reduced pressure. The resulting residue was purified by preparative HPLC (column: Phenomenex luna C18 (250mm × 70mm × 15 um); mobile phase: 40-75% ACN in water (+NH_4_HCO_3_ modifier)) to afford the title compound (480 mg, 26% yield) as a white solid.

^1^H NMR (400 MHz, DMSO-*d*_6_) δ 7.33 - 7.23 (m, 2H), 7.16 (d, *J* = 1.8 Hz, 1H), 6.73 (br t, *J* = 5.2 Hz, 1H), 6.68 (d, *J* = 8.4 Hz, 1H), 6.11 (t, *J* = 6.0 Hz, 1H), 4.01 (d, *J* = 6.0 Hz, 2H), 3.83 (s, 3H), 3.54 - 3.40 (m, 8H), 3.36 (br t, *J* = 5.8 Hz, 4H), 3.31 (s, 2H), 3.05 (q, *J* = 6.0 Hz, 2H), 2.82 (q, *J* = 6.0 Hz, 2H), 2.07 (s, 1H), 1.36 (s, 9H).

LCMS [M+1] = 488.4.

BP1 synthesis, step 4b, *tert*-Butyl N-[2-[2-[2-[2-[[3-methoxy-4-(prop-2-ynylamino)phenyl]sulfonylamino]-ethoxy]ethoxy]ethoxy]ethyl]carbamate: To a solution of *tert*-butyl N-[2-[2-[2-[2-[[3-methoxy-4-(3-trimethylsilylprop-2-ynylamino)phenyl]sulfonylamino]ethoxy]ethoxy]ethoxy]ethyl]-carbamate (580.0 mg, 986.7 μmol, 1.0 equiv.) in MeOH (6.0 mL) was added K_2_CO_3_ (272.7 mg, 1.9 mmol, 2.0 equiv.). The mixture was stirred at 20 °C for 1 hour. The reaction mixture was partitioned between EtOAc (50 mL) and H_2_O (30 mL) and the aqueous layer was extracted with EtOAc (2 × 50 mL). The combined organic layers were washed with brine (50 mL), dried over anhydrous Na_2_SO_4_, filtered, and concentrated under reduced pressure to afford the title compound (400 mg, 79% yield) as an off-white oil.

^1^H NMR (400 MHz, DMSO-*d*_6_) δ 7.34 - 7.24 (m, 2H), 7.15 (d, *J* = 1.8 Hz, 1H), 6.74 (t, *J* = 5.4 Hz, 1H), 6.69 (d, *J* = 8.4 Hz, 1H), 6.13 (t, *J* = 6.0 Hz, 1H), 3.97 (dd, *J* = 2.0, 6.2 Hz, 2H), 3.83 (s, 3H), 3.50 - 3.41 (m, 8H), 3.39 - 3.33 (m, 4H), 3.11 - 3.00 (m, 3H), 2.81 (q, *J* = 6.0 Hz, 2H), 2.07 (s, 3H), 1.36 (s, 9H).

LCMS [M+1] = 416.3.

BP1 synthesis, step 5b, *tert*-Butyl N-[2-[2-[2-[2-[[3-methoxy-4-[3-[4-(tetrahydropyran-4-ylamino)-1-(2,2,2-trifluoroethyl)indol-2-yl]prop-2-ynylamino]phenyl]sulfonylamino]ethoxy]ethoxy]ethoxy]-ethyl]carbamate: To a solution of *tert*-butyl N-[2-[2-[2-[2-[[3-methoxy-4-(prop-2-ynylamino)phenyl]sulfonylamino]ethoxy]-ethoxy]ethoxy]ethyl]carbamate (200 mg, 388 μmol, 1.0 equiv.) in ACN (2.0 mL) was added dichloropalladium-triphenylphosphane (27.2 mg, 38.7 μmol, 0.1 equiv.) and copper(I)iodide (7.4 mg, 38.8 μmol, 0.1 equiv.), 2-iodo-N-tetrahydropyran-4-yl-1-(2,2,2-trifluoroethyl)indol-4-amine (165 mg, 388 μmol, 1.0 equiv.) and triethylamine (162 µL, 1.1 mmol, 3.0 equiv.) under an atmosphere of nitrogen gas. The mixture was stirred at 70°C for 2 hours. The reaction mixture was partitioned between EtOAc (10 mL) and H_2_O (10 mL) and the aqueous layer was extracted with EtOAc (2 × 10 mL). The combined organic layers were washed with brine (50 mL), dried over anhydrous Na_2_SO_4_, filtered, and concentrated under reduced pressure to afford the title compound (200 mg, 64% yield) as a white solid.

^1^H NMR (400 MHz, DMSO-*d*6) δ 7.47 (s, 1H), 7.32 - 7.24 (m, 1H), 7.19 (d, *J* = 1.6 Hz, 2H), 7.08 (s, 1H), 7.00 (t, *J* = 8.0 Hz, 1H), 6.83 (d, *J* = 8.4 Hz, 1H), 6.80 - 6.63 (m, 2H), 6.32 (t, *J* = 6.4 Hz, 1H), 6.21 (d, *J* = 7.8 Hz, 1H), 5.96 - 5.96 (m, 1H), 5.54 (d, *J* = 8.4 Hz, 1H), 5.01 - 4.84 (m, 2H), 4.33 (d, *J* = 6.2 Hz, 1H), 3.93 - 3.83 (m, 4H), 3.51 - 3.39 (m, 9H), 3.38 - 3.34 (m, 1H), 3.38 - 3.29 (m, 7H), 3.04 (q, *J* = 5.8 Hz, 2H), 2.81 (q, *J* = 5.8 Hz, 2H), 2.50 (d, *J* = 1.6 Hz, 72H), 2.04 - 1.97 (m, 1H), 1.91 (br d, *J* = 13.4 Hz, 1H), 1.95 - 1.85 (m, 1H), , 1.36 (s, 9H)

LCMS [M+1] = 812.3.

BP1 synthesis, step 6b, N-[2-[2-[2-(2-aminoethoxy)ethoxy]ethoxy]ethyl]-3-methoxy-4-[3-[4-(tetrahydropyran-4-ylamino)-1-(2,2,2-trifluoroethyl)indol-2-yl]prop-2-ynylamino]benzenesulfonamide: To a solution of *tert*-butyl N-[2-[2-[2-[2-[[3-methoxy-4-[3-[4-(tetrahydropyran-4-ylamino)-1-(2,2,2-trifluoroethyl)indol-2-yl]prop-2-ynylamino]phenyl]sulfonylamino]ethoxy]ethoxy]ethoxy]-ethyl]carbamate (200 mg, 246 μmol, 1.0 equiv.) in DCM (0.6 mL) was added TFA (0.8 mL). The mixture was stirred at 20°C for 1 hour. The reaction mixture was concentrated under reduced pressure to afford the title compound (150 mg, 86% yield) as a brown oil. The crude product was used directly in the next step.

LCMS [M+1] = 712.4.

BP1 synthesis, step 7b, TR-FRET binding probe, BP1, 5-[(3aS,4S,6aR)-2-oxo-1,3,3a,4,6,6a-hexahydrothieno[3,4-d]imidazol-4-yl]-N-[2-[2-[2-[2-[[3-methoxy-4-[3-[4-(tetrahydropyran-4-ylamino)-1-(2,2,2-trifluoroethyl)indol-2-yl]prop-2-ynylamino]phenyl]-sulfonylamino]ethoxy]ethoxy]ethoxy]ethyl]pentanamide: To a solution of N-[2-[2-[2-(2-aminoethoxy)ethoxy]ethoxy]ethyl]-3-methoxy-4-[3-[4-(tetrahydropyran-4-ylamino)-1-(2,2,2-trifluoroethyl)indol-2-yl]prop-2-ynylamino]benzenesulfonamide (40.0 mg, 56.2 μmol, 1.0 equiv.) in DCM (1 mL) was added TEA (28.4 mg, 281 μmol, 39.1 μL, 5.0 equiv.) and (2,5-dioxopyrrolidin-1-yl) 5-[(3aS,4S,6aR)-2-oxo-1,3,3a,4,6,6a-hexahydrothieno[3,4-d]imidazol-4-yl]pentanoate (19.1 mg, 56.2 μmol, 1.0 equiv.). The mixture was stirred at 20°C for 1 hour. The reaction mixture was concentrated under reduced pressure to afford a residue that was purified by preparative HPLC (column: Phenomenex Luna C18 200mm × 40mm × 10um; mobile phase: 35-70% ACN in water (+formic acid modifier)) to afford the title compound (11.2 mg, 21% yield) as a white solid.

^1^H NMR (400 MHz, DMSO-*d*_6_) δ 7.80 (t, *J* = 5.6 Hz, 1H), 7.35 - 7.24 (m, 2H), 7.19 (d, *J* = 1.8 Hz, 1H), 7.08 (s, 1H), 7.01 (t, *J* = 8.0 Hz, 1H), 6.83 (d, *J* = 8.4 Hz, 1H), 6.70 (d, *J* = 8.4 Hz, 1H), 6.40 (s, 1H), 6.36 - 6.28 (m, 2H), 6.22 (d, *J* = 7.8 Hz, 1H), 5.55 (d, *J* = 2.2 Hz, 1H), 4.93 (q, *J* = 9.2 Hz, 2H), 4.41 - 4.22 (m, 3H), 4.18 - 4.07 (m, 1H), 3.95 - 3.79 (m, 5H), 3.52 - 3.36 (m, 14H), 3.23 - 3.13 (m, 2H), 3.11 - 3.02 (m, 1H), 2.90 - 2.74 (m, 3H), 2.57 (d, *J* = 12.6 Hz, 1H), 2.05 (t, J = 7.4 Hz, 2H), 1.91 (d, J = 12.4 Hz, 2H), 1.57 - 1.39 (m, 5H), 1.70 - 1.38 (m, 2H), 1.36 - 1.20 (m, 2H).

LCMS [M+1] = 938.3.
